# Tau Differentially Regulates the Transport of Early Endosomes and Lysosomes

**DOI:** 10.1101/2021.11.01.466759

**Authors:** Linda Balabanian, Dominique V. Lessard, Pamela Yaninska, Piper W. Stevens, Paul W. Wiseman, Christopher L. Berger, Adam G. Hendricks

## Abstract

Microtubule-associated proteins (MAPs) modulate the motility of kinesin and dynein along microtubules to control the transport of vesicles and organelles. The neuronal MAP tau inhibits kinesin-dependent transport. Phosphorylation of tau at tyrosine 18 by fyn kinase results in weakened inhibition of kinesin-1. We examined the motility of early endosomes and lysosomes in cells expressing wild-type (WT) tau and phosphomimetic Y18E tau. Lysosome motility is strongly inhibited by tau. Y18E tau preferentially inhibits lysosomes in the cell periphery, while centrally located lysosomes are less affected. Early endosomes are more sensitive to tau than lysosomes, and are inhibited by both WT and Y18E tau. Our results show that different cargoes have disparate responses to tau, likely governed by the types of kinesin motors driving their transport. In support of this model, kinesin-1 and -3 are strongly inhibited by tau while kinesin-2 and dynein are less affected. In contrast to kinesin-1, we find that kinesin-3 is strongly inhibited by phosphorylated tau.

## Introduction

A web of interactions between motor proteins, cargo adapters, and the cytoskeleton governs the transport of vesicles and organelles. The microtubule tracks control intracellular transport through microtubule-associated proteins (MAPs), tubulin post-translational modifications, and the organization of the microtubule network. Teams of kinesin and dynein motor proteins associate with cargoes, which confer bidirectional motility and the flexibility to navigate obstacles and readily change microtubule tracks. Cytoplasmic dynein is responsible for retrograde transport in animal cells, while kinesins-1, -2, -3 and -4, drive anterograde transport (Balabanian et al., 2018; Ghiretti et al., 2016; Hancock, 2014; Hirokawa and Takemura, 2005; Janke, 2014; Verhey and Gaertig, 2007).

The motility and positioning of lysosomes, driven by dynein and kinesins 1, 2, and 3, are crucial for robust autophagy and immune responses (Pu et al., 2016). Kinesin-2 is required for the transport of late endosomes and lysosomes, but not early endosomes (Brown et al., 2005). In addition to kinesin-2, subpopulations of lysosomes are driven alternatively by kinesin-1 on juxtanuclear acetylated microtubules and kinesin-3 (KIF1Bβ and KIF1A) on peripheral tyrosinated microtubules (Bentley et al., 2015; Guardia et al., 2016; Matsushita et al., 2004; Norris et al., 2014). Early endosomes, responsible for sorting endocytosed particles for recycling or lysosome-mediated degradation, are driven in the anterograde direction by kinesin-1, as well as members of the kinesin-3 family (KIF16B, KIF13A, KIF13B) (Bentley et al., 2015; Blatner et al., 2007; Delevoye et al., 2014; Hoepfner et al., 2005). Recent studies from multiple groups indicate that kinesin-3 motors conduct transport of various cargoes on dynamic microtubules in more peripheral regions and presynaptic sites in neurons, whereas kinesin-1 acts on stable acetylated microtubules closer to the nucleus (Cai et al., 2009; Guedes-Dias et al., 2019; Katrukha et al., 2017; Konishi and Setou, 2009; Serra-Marques et al., 2020; Tas et al., 2017).

The MAP tau stabilizes and bundles microtubules, acting as a spacer between neighbouring microtubules in bundles (Kanai et al., 1992; Scott et al., 1992; Takemura et al., 1992). Tau also inhibits the motility of motor proteins by blocking their interaction with microtubules (Dixit et al., 2008; Hoeprich et al., 2014; McVicker et al., 2011; Monroy et al., 2018; Vershinin et al., 2007). Axonal microtubule bundles are heavily decorated with tau, yet robust intracellular transport in axons supports neuronal signalling and homeostasis (Kanai and Hirokawa, 1995; Kosik and Finch, 1987). Different types of kinesin are differentially affected by the presence of MAPs on the microtubule surface. Kinesin-1 does not readily circumvent tau obstacles on microtubules, whereas kinesin-2 and kinesin-8 navigate around tau (Guzik-Lendrum et al., 2015; Hoeprich et al., 2014; Malaby et al., 2019; Shastry and Hancock, 2011).

Tau rapidly switches between states that statically bind or dynamically diffuse over the microtubule surface (Hinrichs et al., 2012; McVicker et al., 2014). The shortest tau isoform of the central nervous system, tau-3RS, binds more statically to the microtubule and most strongly inhibits kinesin *in vitro* (Dixit et al., 2008; McVicker et al., 2011; Vershinin et al., 2007). Post-translational modifications such as phosphorylation alter tau’s conformation and surface charge, modulating its microtubule binding. Phosphomimetic tau (Y18E) has reduced affinity on reconstituted microtubules, increasing its diffusivity and consequently alleviating tau’s inhibitory effect on purified kinesin-1 motors (Stern et al., 2017). As such, the phosphorylation state of tau is an important factor in determining the processivity of motor motility.

Over 85 putative phosphorylation sites have been identified on tau, phosphorylated by multiple kinases including GSK3ß, AKT, MAPK, and Fyn (Sergeant et al., 2008). Phosphorylation of tau at multiple sites is prevalent in tauopathies and the aggregation of the hyper-phosphorylated misfolded tau is correlated with worsening cognitive impairment (Ballatore et al., 2007). The nonreceptor tyrosine kinase fyn phosphorylates tau at Y18 (Lee et al., 2004), the last residue of the phosphatase-activating domain (PAD) of tau. Y18 phosphorylation prevents the PAD from activating the protein phosphatase 1 (PP1) and GSK-3 pathways, which otherwise inhibit anterograde fast axonal transport and mediate kinesin detachment from cargoes. Subsequently, Y18 phosphorylation of tau counteracts the inhibition by tau aggregates on anterograde fast axonal transport (Kanaan et al., 2012; Kanaan et al., 2011; LaPointe et al., 2009; Morfini et al., 2004). Furthermore, PAD exposure and activity in tau aggregates found in Alzheimer’s disease precede the phosphorylation at Y18 in aggregates during disease progression (Kanaan et al., 2012). As such, fyn kinase-mediated phosphorylation at Y18 may confer a protective effect against tau inhibition of cargo transport.

Motivated by the apparent paradox between the observations that tau inhibits kinesin motility yet robust intracellular transport occurs along axonal microtubules decorated by tau, we asked if tau phosphorylation at Y18 might modulate tau’s inhibition of organelle transport in cells. Further, we asked if the effect of phosphorylation is different for cargoes driven by different types of kinesin motors (kinesin-1, -2, and -3), given the varying sensitivity of motors to tau *in vitro*. We studied the localization and movement of lysosomes and early endosomes in cells exogenously expressing wild-type (WT) tau (shortest isoform tau-3RS) alone or with fyn kinase, or phosphomimetic tau (Y18E). We accounted for variable expression levels of tau in cells by quantifying tau intensity and measuring the response in the motility of cargoes as a function of the level of tau.

At low levels of tau expression, we found that phosphomimetic and phosphorylated tau partially rescue the inhibition of lysosome motility, compared to WT tau. Tau inhibition is specifically reduced for the subpopulation of lysosomes positioned in the perinuclear or juxtanuclear region, which are expected to be transported on stable acetylated microtubules primarily by kinesin-1 (Cai et al., 2009; Katrukha et al., 2017). In contrast, peripheral lysosomes, which localize on dynamic tyrosinated microtubules by kinesin-3 (Guardia et al., 2016; Guedes-Dias et al., 2019; Konishi and Setou, 2009; Norris et al., 2014; Serra-Marques et al., 2020; Tas et al., 2017), were inhibited by phosphomimetic tau. WT tau inhibits both juxtanuclear and peripheral sub-populations of lysosomes similarly. Unlike lysosomes, early endosomes are inhibited by both phosphomimetic/ phosphorylated tau and unphosphorylated (WT) tau. Using *in vitro* motility assays, we show that kinesin-3 motor proteins are inhibited by both phosphomimetic tau and WT tau on reconstituted, paclitaxel-stabilized microtubules. Previously, we showed that phosphomimetic tau, unlike WT tau, does not strongly inhibit kinesin-1 (Stern et al., 2017). Microtubule bundling by tau is also expected to affect cargo transport. We observe that global rearrangements of the microtubule network occurred only with high levels of tau. Together, our results show that regulation of tau by Y18 phosphorylation differentially affects the displacement of specific cargoes.

## Results

### Tau biases lysosome motility and localization to the perinuclear region

We examined the directionality of lysosomes in control COS-7 fibroblast cells, and the localization of tau in cells transfected with WT tau (wild-type) or phosphomimetic tau (Y18E) (Fig. 1A, B). COS-7 do not endogenously express tau, such that the type of tau present can be tightly controlled. The flat morphology of COS-7 cells allows for high-resolution imaging using TIRF (Total Internal Reflection Fluorescence) microscopy (Movie S1). Lysosomes are localized throughout the cell and concentrated in the perinuclear region, close to the microtubule-organizing center (MTOC) (Fig. 1A, B) (Pu et al., 2016). Around half of all lysosomes are stationary in control cells (∼51%), and the remaining lysosomes are evenly divided between active transport towards and away from the cell center (Fig. 1C, D).

**Figure 1:**
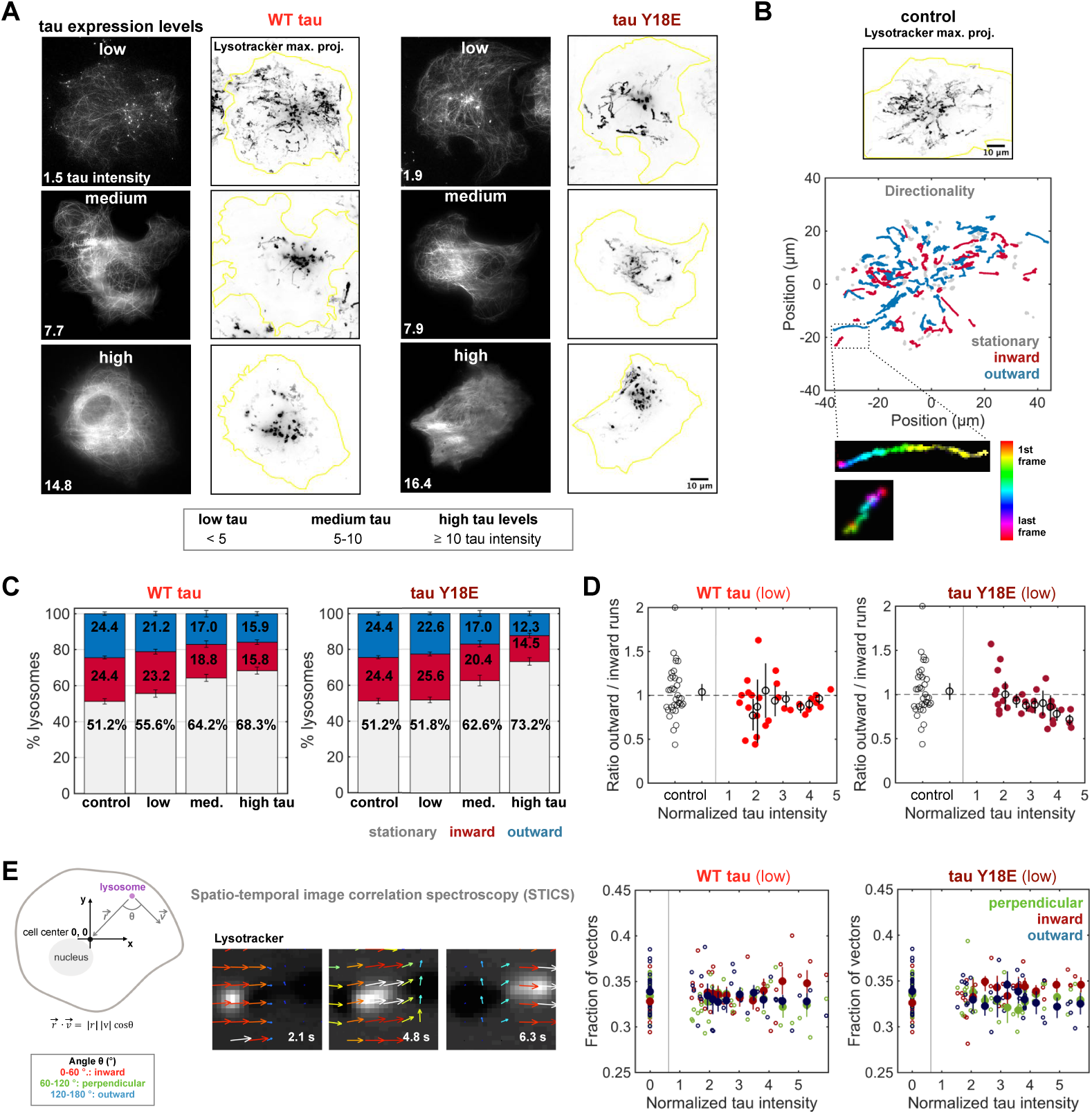
Tau causes retrograde bias of lysosome motility. **A**. Transient transfection results in variable tau-mApple expression levels. In cells expressing low levels (normalized tau intensity < 5), tau enriches on microtubules and lysosomes are distributed throughout the cell, as seen on the maximum projections of lysosome movies. In cells expressing medium (5 ≤ normalized tau intensity < 10) and high (normalized tau intensity ≥ 10) levels of tau, tau is localized both along microtubules and in the cytosol. Lysosomes are more constrained to the perinuclear region in these cells. **B**. The maximum projection image show lysosomes exhibit robust motility along microtubules in a control cell (with no tau expression). The net directionality of lysosome trajectories, tracked with Trackmate (Tinevez et al., 2016), was categorized as inward (towards cell center) or outward (towards the cell periphery) for motile lysosomes (with radius of gyration Rg ≥ 0.5 μm), or as stationary (Rg < 0.5 μm). **C**. The fraction of stationary lysosomes increases with the level of tau expression (mean ± SEM). **D**. The fraction of inward and outward trajectories is approximately equal for motile lysosomes in control cells. The ratio of the number of outward to inward indicates that WT and Y18E tau reduce the fraction of outward movement. Small symbols indicate single cells, large symbols indicate the means ± 95% confidence intervals (error bars) calculated using bootstrapping. **E**. Spatiotemporal Image Correlation Spectroscopy (STICS) was used to calculate velocity fields of the lysosome movement. θ is defined as the angle between the velocity (v) vector and the vector (r) pointing from the lysosome to the cell center (0, 0). An angle of 0-60° indicates inward movement (toward cell center), 60-120° indicates perpendicular movement with respect to the cell center (depicted on the schematic), and 120-180° represents outward motion. The fraction of lysosome trajectories moving inward, outward or perpendicular in respect to the cell center (for each cell) shows lysosomes move inward more often in cells expressing low levels of WT tau or Y18E tau, relative to control cells (small symbols: single cell, large symbols: mean ± 95% confidence intervals calculated using bootstrapping). Both the tracking-based directionality analysis (in C, D) and the STICS analysis (in E) demonstrate that lysosomes exhibit an inward bias in response to tau. The plots for the datasets showing the cells over the whole range of tau expression levels (low, medium, high) are found in Fig. S1D, E. (Control: n = 35 cells in C, D and 33 in E; WT tau: 71; Y18E tau: 83 cells.)

We quantified the levels of tau expression in transfected cells by measuring tau-mApple intensity and normalizing to the background intensity outside of the cell. Tau enriches strongly on microtubules, with low cytosolic signal present in cells with low levels of tau (normalized tau intensity < 5) (Fig. 1A, 1st row). Cells expressing medium levels (5 ≤ normalized tau intensity < 10) show a considerable cytosolic signal in the perinuclear region (Fig. 1A, 2nd row) and finally high levels of tau expression (normalized tau intensity ≥ 10) exhibit cell-wide cytosolic signal (Fig. 1A, 3rd row). We also co-expressed WT tau with fyn kinase (Fig. S1A-E), which phosphorylates tau at Y18 (Fig. S1F) (Lee et al., 2004), and found that fyn alone influences lysosome movement, thus we focused on directly comparing WT tau and tau Y18E. Lysosomes are more constrained to the perinuclear region (Fig. 1A) and the percentage of stationary lysosomes accumulates with increasing tau expression, notably with medium and high levels of tau (Fig. 1C, Fig. S1D), such that it distorts the analysis on the effects of tau on lysosome directionality in these cells (Fig. S1D, E). Low levels of tau in cells are comparable with endogenous tau levels in neuronal axons (Fig. S1F-H) (Black et al., 1996; Xia et al., 2016). Consequently, we focus our analysis to cells expressing low levels of WT tau and phosphomimetic Y18E tau.

Microtubules exhibit robust dynamics in cells expressing low levels of WT tau or Y18E tau (Fig. S2A). However, microtubule organization is altered dramatically with high levels of tau expression (Fig. S2). Tau binds along the length of microtubules (Fig. S2B) (Samsonov et al., 2004), and bundles microtubules, acting as a spacer between adjacent microtubules (Chen et al., 1992; Kanai et al., 1992; Scott et al., 1992; Takemura et al., 1992). We studied peripheral microtubule density and bundling in cells to distinguish the effect of tau on cargo transport from its indirect influence on cargo motion through remodelling of the microtubule network (Fig. S2). High levels of tau dramatically rearrange the microtubule network into thick microtubules bundles, in a ring-shaped array in the periphery (Fig. S2C-F) (Yu et al., 2016). Lysosomes are trapped within the bundled microtubule rings and some lysosomes are seen circling around the interior edges (Fig. S2G). Fyn kinase overexpression alone also increases peripheral microtubule density (Fig. S2C-F). At low levels of WT tau or tau Y18E expression, most cells retained a radial organization of microtubules similar to control (Fig. S2E). Overall, lysosome directionality (Fig. 1), motility (Fig. 2, Fig. 3) and localization (Fig. 3) are unlikely to be affected by tau’s effect on microtubule organization at low tau levels, as extreme reconfigurations of the microtubule network occur with high amounts of tau.

**Figure 2:**
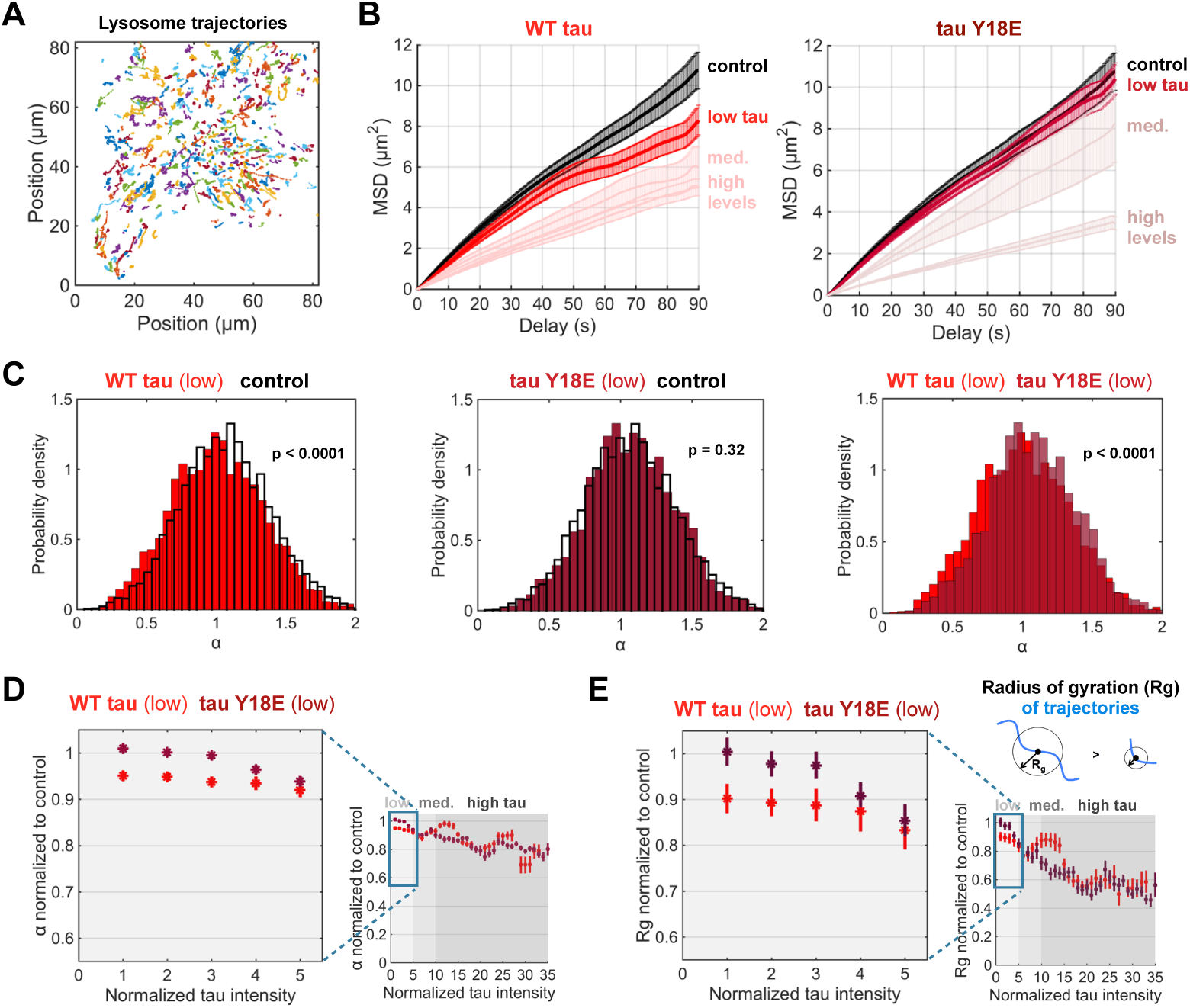
Phosphorylation of tau at Y18 relieves inhibition of lysosome motility. **A**. Lysosomes were imaged for 90 seconds and tracked using Trackmate in ImageJ (Tinevez et al., 2016). Lysosome trajectories from a control cell are shown here. **B**. The mean-squared displacement (MSD) of lysosomes in cells expressing low levels of tau Y18E are similar to control, whereas low levels of WT tau inhibit lysosome motility (mean ± SEM). **C**. The slope of the MSD (α) indicates the proportion of stationary (α=0), diffusive (α=1), and processive (α=2) motility. α values for cells expressing low levels of WT tau are reduced compared to control and tau Y18E (1-way ANOVA, Tukey’s test, p < 0.0001). Low levels of tau Y18E do not decrease lysosome processivity compared to control (p = 0.32). **D**. Sliding means of the α values were calculated as a function of tau intensity for trajectories in cells expressing low levels of WT tau or tau Y18E (left plot). Processivity (α) decreases progressively with tau intensity (right plot, mean ± 95% confidence intervals). Control: n = 4882 trajectories; fyn: 2149; WT tau: 7910; Y18E tau: 9970; WT tau + fyn: 4644 trajectories. **E**. The radius of gyration (Rg) is an indicator of the mean distance traveled in a trajectory (see Methods), where a circle with radius Rg contains half of the spots of the trajectory. Lysosome displacement is inhibited less by phosphomimetic Y18E tau than WT tau (left plot). The displacement of lysosomes decreases with increasing tau intensity, over the wider range of tau intensities (right plot, mean ± 95% confidence intervals). Control: n = 5408 trajectories, from n = 35 cells, over 20 experiments; WT tau: 9024 traj., 73 cells, over 14 experiments; Y18E tau: 11105 trajectories, 83 cells, over 5 experiments. This analysis uses the same data set as in Fig. 1.

**Figure 3:**
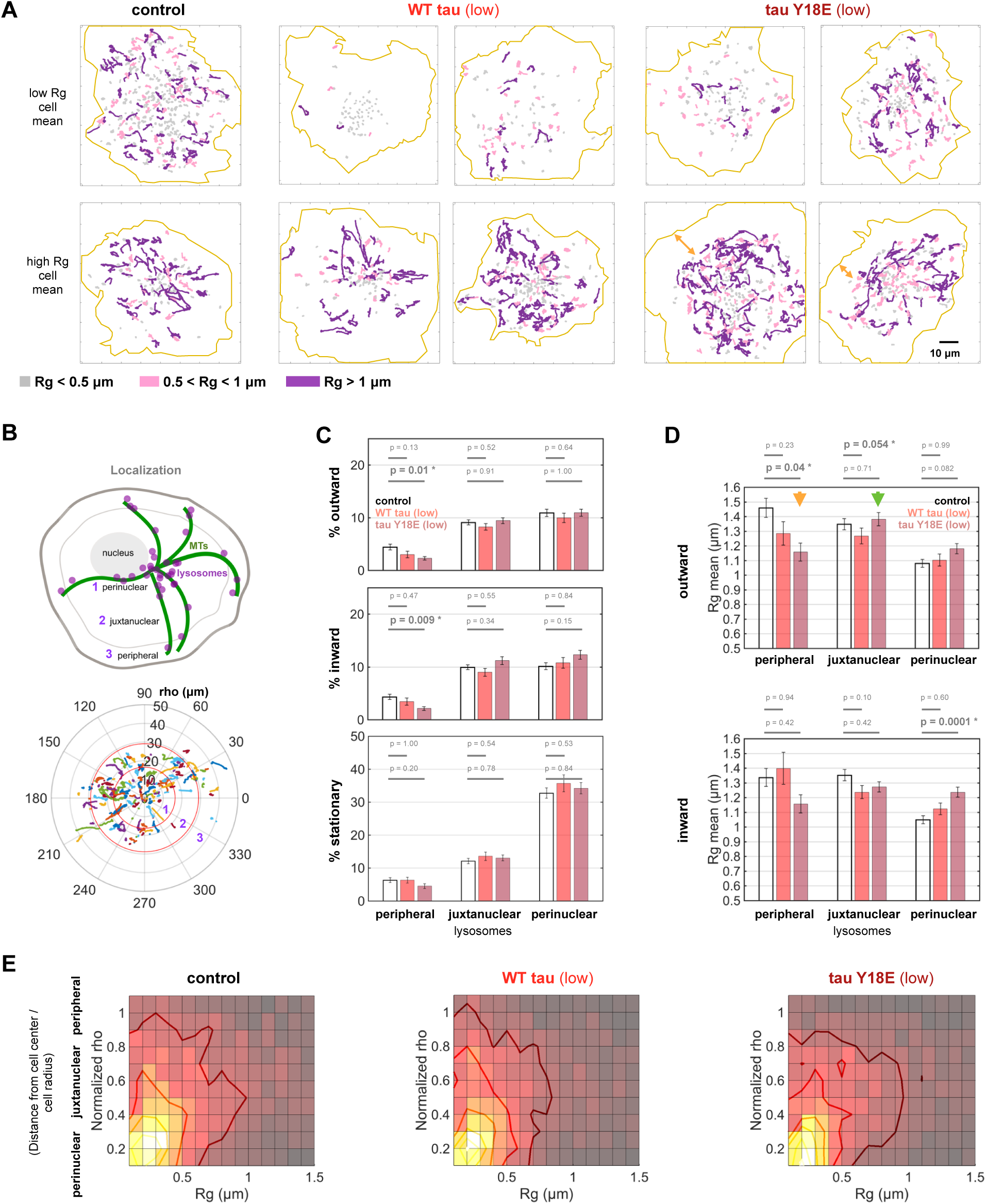
Phosphomimetic tau Y18E inhibits the motility of lysosomes in the cell periphery. **A**. Stationary (gray), short (magenta), and long (purple) trajectories (over 90 sec. movies) are distributed throughout the control cells. WT tau expression results in fewer long trajectories in all regions of the cell. In contrast, Y18E tau expression preferentially inhibits long trajectories in the cell periphery and often precludes lysosomes in this region (orange arrows). For comparison, we show representative images for cells with low and high average lysosome displacement (Rg). **B**. Rho is defined as the mean distance of each trajectory from the cell center, normalized to the average radius of the cell. Lysosomes were categorized as perinuclear (rho < 0.5), juxtanuclear (0.5 ≤ rho < 0.85), and peripheral (rho ≥ 0.85). **C**. The percentage of inward-directed, outward-directed, and stationary lysosomes in different regions of the cell (mean ± SEM) indicates the proportion of moving lysosomes in both the inward and outward directions is significantly reduced in the periphery of cells expressing Y18E tau. **D**. The mean Rg (± SEM) of motile lysosomes (Rg > 0.5 μm) moving in the outward (upper panels) or inward direction (bottom panels) is shown for each region of the cell (peripheral, juxtanuclear or perinuclear localization). Juxtanuclear lysosomes maintain outward displacement similar to control in the presence of Y18E tau (green arrow), while peripheral lysosomes are inhibited by tau Y18E (orange arrow). p-values from 1-way ANOVA and Tukey’s test are shown on bar charts for low levels of WT tau or Y18E tau expression compared to control. **E**. The distribution of lysosomes with respect to their Rg and localization. The distribution of moving lysosomes (Rg ≥ 0.5 μm) in Y18E tau expressing cells is concentrated in the juxtanuclear region while moving lysosomes are more evenly distributed across regions in control and WT tau expressing cells. (Control: n = 35 cells, WT tau (low): 29 cells; Y18E tau (low): 31 cells). This analysis uses the same data set as in Fig. 1.

Both low levels of WT tau and phosphomimetic Y18E tau cause an increase of lysosome motility towards cell center, as shown by tracking analysis (Fig. 1D, 95% confidence intervals). Single-particle tracking can be challenging in situations where particles are clustered. Thus, we applied Spatio-temporal image correlation spectroscopy (STICS) which generates vector fields based on the correlation of pixel intensities over a range of spatial and temporal scales (Fig. 1E, Fig. S1E). STICS also demonstrates that tau results in an inward bias in lysosome motility (Fig. 1E, 95% confidence intervals). Both single-particle tracking and correlation analysis show that tau biases lysosome motility towards the cell center, in agreement with *in vitro* motility assays of isolated phagosomes (Chaudhary et al., 2018), and leads to the enrichment of stationary lysosomes in the perinuclear region with increasing levels of tau.

### Phosphorylation at Y18 reduces tau-mediated inhibition of lysosome transport

Using high-resolution tracking, we analyzed the motility of lysosomes stained with Lysotracker Deep Red in live cells (Fig. 2A). We measured the mean-squared displacement (MSD) of lysosome trajectories and calculated the slope (α value) to determine the type of motion (α < 1 for confined, α = 1 for diffusive, α > 1 for directed motion)(Tarantino et al., 2014). We also calculated the radius of gyration (Rg) of each trajectory as a measure of the distances travelled by the cargoes. MSD analysis shows that low levels of WT tau constrain lysosomal motion, while lysosome processivity in cells expressing phosphomimetic Y18E tau remains similar to control cells (Fig. 2B). The processivity (α) of lysosomal trajectories decreases in cells expressing WT tau compared to control (1-way ANOVA, Tukey’s test, p < 0.0001) and compared to tau Y18E (p < 0.0001), while the distributions remains similar for control cells and tau Y18E cells (p = 0.32)(Fig. 2C). Overall, the motility of lysosomes decreases with increasing levels of tau (Fig. 2D, E, Fig. S3D, E, 95% confidence intervals). WT tau reduces processivity (Fig. 2D) and distances travelled by lysosomes (Fig. 2E), even at low levels of tau expression. Interestingly, at low levels, phosphomimetic Y18E tau does not decrease the processivity and displacement of lysosomes, which remain similar to control (Fig. 2D, E, 95% confidence intervals).

Fyn kinase decreases lysosomal displacement and processivity. However, co-expression of fyn and low levels of WT tau did not lead to a further reduction, suggesting that phosphorylated tau does not strongly inhibit lysosome movement (Fig. S3A-C). The α distribution for cells expressing fyn alone (2-way ANOVA, p = 0.0008) or with WT tau also demonstrates that phosphorylated tau (WT tau + fyn) does not behave like WT tau (interaction tau and fyn: p = 0.013) (Fig. S3B). Together, these results show that low levels of phosphomimetic or phosphorylated tau at Y18 do not strongly inhibit lysosome motility, compared to non-phosphorylated (WT) tau.

We examined the speed and pausing behaviour of lysosomes (Fig. S3F-H). Lysosome trajectories typically consist of motile and paused segments (Fig. S3F, G). The average speed of lysosomes when excluding paused segments does not vary considerably with increasing levels of tau expression (Fig. S3H). Comparatively, the average speed when including detected paused segments of trajectories decreases to a greater extent, as the percentage of time spent pausing within trajectories increases with tau (Fig. S3H). At low levels of expression, lysosomes in phosphomimetic tau Y18E expressing cells exhibited a lower average speed and higher percentage of time spent pausing relative to WT tau (Fig. S3H). Taken together with the finding that lysosomes travel longer distances in the presence of phosphomimetic tau, these results suggest that lysosomes are more likely to continue moving after a pause in the presence of tau phosphorylated at Y18.

### Phosphomimetic tau (Y18E) inhibits the motility of peripheral lysosomes but not juxtanuclear lysosomes

The distribution of both stationary lysosomes and lysosomes moving longer distances is dispersed uniformly in perinuclear, juxtanuclear, and peripheral regions of control cells and cells expressing low levels of WT tau (Fig. 3A). In contrast, lysosomes with long runs (Rg ≥ 1 μm) are concentrated in the juxtanuclear region of tau Y18E expressing cells (Fig. 3A). Relatively fewer lysosomes localized in the periphery with phosphomimetic Y18E tau (Fig. S4A). Stationary lysosomes or lysosomes with short runs are still found in the periphery of some cells (Fig 3A top panels) and nearly absent altogether in others (Fig. 3A bottom panels, orange arrows). With phosphomimetic Y18E tau, cells which have fewer peripheral lysosomes (Fig. 3A bottom panels, Fig. S4B) show a higher mean for lysosome displacement (Spearman’s correlation coefficient ρ = -0.41, p = 0.02), while no correlation is found for control and WT tau expressing cells (Fig. S4C). Phosphomimetic Y18E tau expression, unlike WT tau, is differentially affecting the motility of lysosomes positioned centrally or peripherally.

Lysosome trajectories were projected on polar plots to measure their distance from cell center and localization was categorized into three sub-cellular regions: peripheral, juxtanuclear and perinuclear (Fig. 3B, see Methods). Cells expressing even low levels of phosphomimetic Y18E tau show a significant loss of outward and inward-moving lysosomes in the periphery compared to control (1-way ANOVA, Tukey’s test, p = 0.01 and p = 0.009 respectively), which is not seen with low levels of WT tau (Fig. 3C). The loss of peripheral lysosomes occurs more prominently with tau Y18E than with unphosphorylated (WT) tau (Fig. 3A, C, Fig. S4A).

Overall, the localization of lysosomes in the periphery and the motility of lysosomes drops gradually with increasing tau levels, consistent with inhibition of kinesin-dependent transport towards distal regions of the cell (Fig. S4D, E). Previous studies suggest that centrally localized lysosomes are to a large extent driven by kinesin-1 on acetylated microtubules, whereas peripheral lysosomes are transported primarily by kinesin-3 on tyrosinated microtubules (Guardia et al., 2016; Serra-Marques et al., 2020; Tas et al., 2017). As such, we hypothesized that juxtanuclear and peripheral lysosomes would be affected to different extents by tau. In cells expressing low levels of WT tau, lysosomes positioned peripherally or in the juxtanuclear region are inhibited (log-normalization, 1-way ANOVA, Tukey’s test, p = 0.054) and exhibit similar displacement in the outward direction (Fig. 3D). However, in cells expressing low levels of phosphomimetic Y18E tau, juxtanuclear lysosomes travel distances similar to control (green arrow, p = 0.71), while motility is reduced for peripheral lysosomes (Fig. 3D, orange arrow, p = 0.04). As expected, low levels of WT tau did not inhibit inward-moving peripheral lysosomes (Fig. 3D, p = 0.94), as the dynein motors responsible for inward transport are less sensitive to tau (Chaudhary et al., 2018; Dixit et al., 2008; Vershinin et al., 2008). Y18E tau increases the displacement of lysosomes in the perinuclear region (Fig. 3D, p = 0.0001), suggesting the motors that transport lysosomes near the cell center are not inhibited by Y18E tau. STICS analysis reveals an inward bias in the movement of remaining lysosomes in the periphery (Fig. S4F), which could lead to the overall loss of peripheral lysosomes over time with Y18E tau, coincidentally where acetylated microtubules are scarce (Fig. S4G). The persistent motility of lysosomes in Y18E tau cells in central regions corresponds with the regions enriched in acetylated microtubules (Fig. S4G).

Fyn kinase increases the fraction of stationary lysosomes in the periphery and juxtanuclear region (Fig. S4A, Fig. S4D, 2-way ANOVA, p = 0.013 and p = 0.0028 respectively) and reduces the fraction of perinuclear moving lysosomes (Fig. S4D, p = 0.0013 and p = 0.0016 for outward and inward-moving lysosomes respectively). Fyn is a non-receptor tyrosine kinase associated with membrane components and involved in many signalling pathways controlling morphological differentiation and cytoskeletal rearrangements near the membrane for process outgrowth (Klein et al., 2002; White and Kramer-Albers, 2014), which could explain why microtubule density in the periphery is also altered with fyn over-expression (Fig. S2C-F) and why lysosomes are more enriched in the periphery of fyn-transfected cells (Fig. S4A, D). Most interestingly, we show that motile lysosomes (Rg ≥ 0.5 μm) are more enriched in the juxtanuclear region of cells expressing low levels of phosphomimetic Y18E tau, compared to the more dispersed distribution of lysosomes found in control cells and cells expressing low levels of WT tau (Fig. 3E).

### Early endosome motility is inhibited by both WT tau and Y18 phosphorylated tau

The observation that sub-populations of lysosomes positioned centrally or peripherally were disparately affected by tau suggested that tau phosphorylation might differentially affect other organelles. Streptavidin quantum dots (Qdots) coated with biotinylated epidermal growth factor (EGF) were endocytosed into COS-7 fibroblast cells to label early endosomes (Zajac et al., 2013)(Fig. 4A). Qdot-containing early endosomes exhibit shorter bursts of directed motion (Fig. 4A, Fig. S5A, B, Movie S2), comparatively to lysosomes which are more processive and bidirectional (Fig. 2A). Fyn kinase over-expression highly inhibits early endosome motion with or without tau (Fig. S5A-D). Directed motion is still observed with low levels of WT tau or phosphomimetic Y18E tau, but nearly absent for cells with medium and high levels of tau (Fig. S5B, E, F).

**Figure 4:**
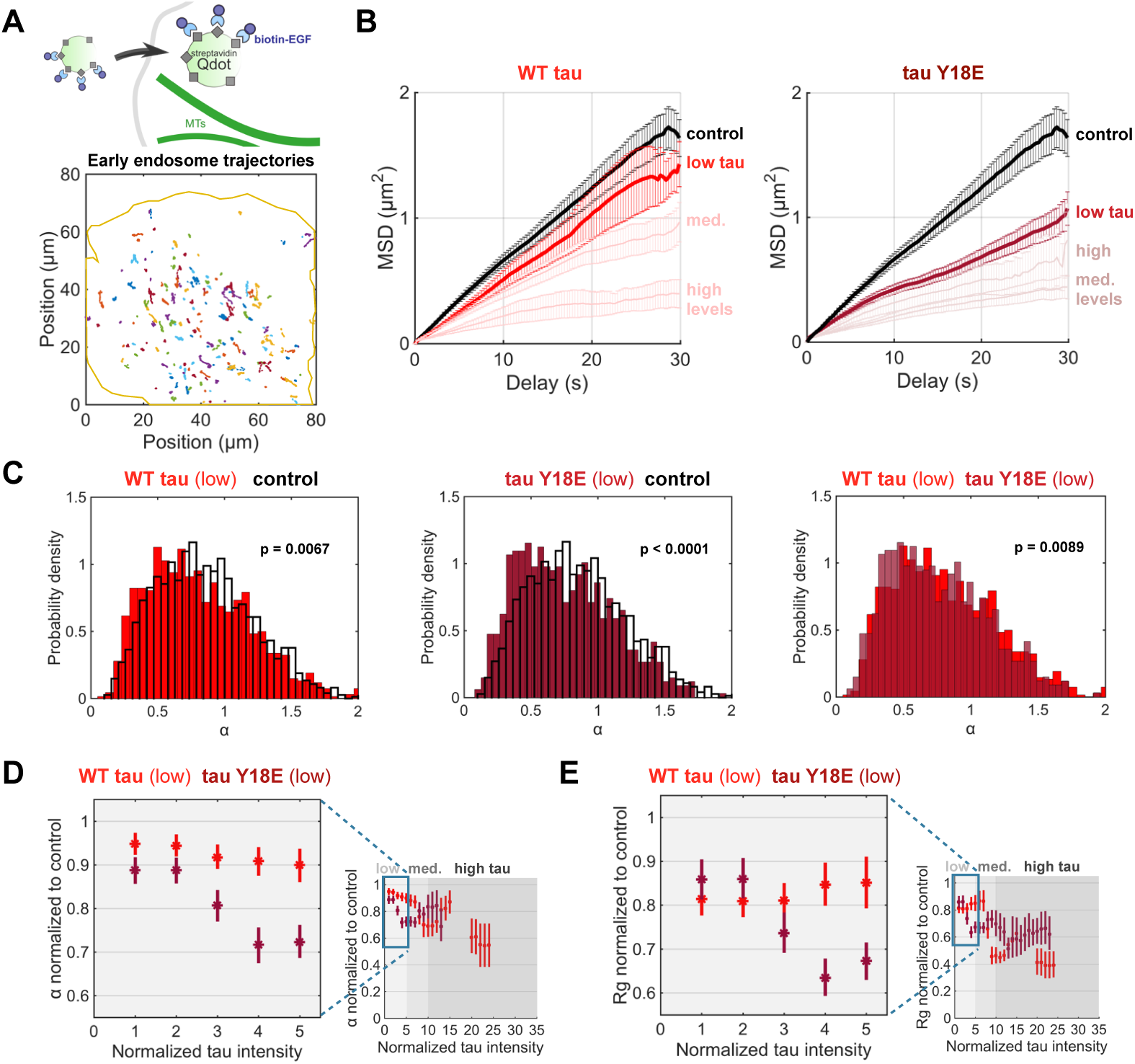
WT and phosphomimetic tau strongly inhibit early endosome motility. **A**. EGF- coated Qdots were used to image early endosomes (< 1 hour post-internalization) and tracked using Trackmate (Tinevez et al., 2016). Early endosomes exhibit short processive runs (control cell shown here). **B**. MSD analysis shows that both low levels of WT tau and Y18E tau inhibit early endosome motility. In contrast to lysosomes, early endosome motility is more strongly inhibited by Y18E tau than WT tau. **C**. The distribution of MSD α values indicates that WT and Y18E tau reduce early endosome processivity (1-way ANOVA, Tukey’s test). **D**. The MSD α (mean ± 95% confidence intervals) of early endosomes, normalized to the mean α of control trajectories, shows inhibition of early endosome processivity with low levels of WT tau and Y18E tau. Control: n = 1394 trajectories; fyn: 148; WT tau: 1653; Y18E tau: 1185; WT tau + fyn: 338 trajectories. **E**. The radius of gyration (Rg) of early endosome trajectories similarly shows early endosome inhibition by both WT tau and Y18E tau. Thus, Y18 phosphorylation of tau does not relieve tau-mediated inhibition of early endosomes. (Control: n = 1946 trajectories, from 28 cells, over 19 experiments; WT tau: 2783 traj., 84 cells, over 15 experiments; Y18E tau: 2193 trajectories, 53 cells, over 6 experiments.)

MSD analysis shows that both low levels of WT tau and tau Y18E inhibit early endosome motility, and that phosphomimetic Y18E tau is more inhibitory to these cargoes than WT tau (Fig. 4B) The processivity (α) of trajectories in cells expressing low levels of WT tau and phosphomimetic Y18E tau is reduced relative to control (1-way ANOVA, Tukey’s test, p = 0.0067 and p < 0.0001 respectively) (Fig. 4C). Analysis of trajectories of Qdot-containing early endosomes revealed that WT tau and phosphomimetic tau both inhibit the processivity (Fig. 4D) and displacement (Fig. 4E, 95% confidence intervals) of early endosomes. STICS analysis shows that the fraction of outward movement of early endosomes is reduced in Y18E tau-expressing cells (Fig. S5G). In addition, early endosomes are more sensitive to tau compared to lysosomes as their displacement is highly inhibited at even medium levels of tau, while lysosomes reached that level of inhibition with higher amounts of tau (Fig. 2D, E, Fig. 4D, E). All in all, phosphorylation of tau at Y18 does not counteract the inhibitory influence of tau on early endosomes.

### WT tau and phosphomimetic tau at Y18 inhibit kinesin-3 *in vitro*

Peripheral lysosomes and early endosomes show reduced motility in the presence of low levels of phosphomimetic Y18E tau, while juxtanuclear lysosomes demonstrate motility similar to control. Kinesin-3 contributes to the motility of the cargoes that are sensitive to Y18E tau (Bentley et al., 2015; Guardia et al., 2016; Hoepfner et al., 2005), leading to the hypothesis that kinesin-3, unlike kinesin-1, is inhibited by phosphomimetic tau. To investigate the effect of phosphomimetic tau on kinesin-3 directly, we performed kinesin-3 (KIF1A) motility assays on reconstituted microtubules *in vitro* (Fig. 5A). KIF1A run lengths dropped to nearly half of that of control with 200 nM WT tau (equivalent to 1:5 tau:tubulin ratio)(1-way ANOVA, Tukey’s test, p < 0.0001), and were reduced even further with phosphomimetic Y18E tau (Fig. 5B, p < 0.0001). The overall speeds of the motor protein is relatively higher with WT tau compared to control due to reduced inter-run pausing (Fig. 5C, p = 0.0002, Fig. 5A). We previously showed that kinesin-1 run lengths are also strongly reduced by WT tau, but that the inhibition was relieved with phosphomimetic Y18E tau (Stern et al., 2017)(Fig. 5D). Thus, kinesin-1 and kinesin-3 are differentially affected by tau phosphorylation at Y18 (Fig. 5D).

**Figure 5:**
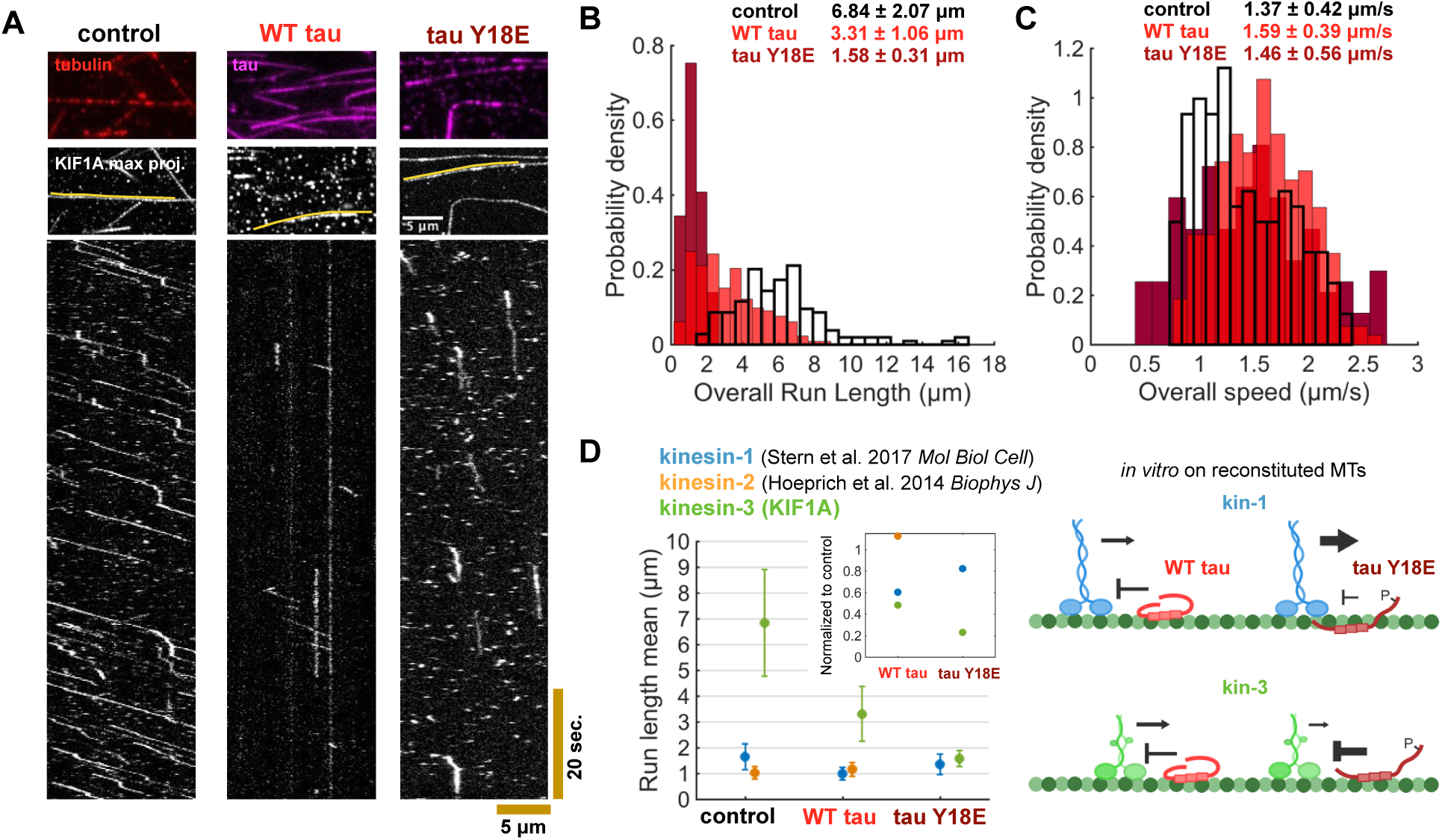
Kinesin-3 (KIF1A) is inhibited by WT and phosphomimetic Y18E tau *in vitro*. **A**. Single molecule motility assays (Movie S3) on reconstituted microtubules indicate that KIF1A(1-393)-LZ-3xmCitrine processivity is strongly reduced in the presence of WT and Y18E tau, compared to control (no tau), as seen on kymographs. **B**. WT tau and tau Y18E decrease KIF1A run lengths (1-way ANOVA, Tukey’s test, p < 0.0001), with Y18E tau causing a greater inhibition than WT tau (p < 0.0001). **C**. KIF1A exhibits fewer inter-run pauses (A) in the presence of WT tau, resulting in higher velocities (p = 0.0002). Y18E tau’s effect on KIF1A’s speed is not significant (p = 0.35). Control: 144 trajectories, WT tau: 212, Y18E tau: 153 trajectories. **D**. Tau phosphorylation has differential effects on kinesin-1 (Stern et al., 2017) and kinesin-3 (as seen in A-C) on reconstituted microtubules (MTs) *in vitro*. Experimental conditions were similar (kinesin-1 and kinesin-2: 10 mM PIPES, 50 mM potassium acetate, 4 mM magnesium acetate; kinesin-3: 12 mm PIPES, 1 mm MgCl_2_; All: supplemented with 1 mM EGTA, 10 mM DTT, and an oxygen scavenger system (5.8 mg/mL glucose, 0.045 mg/mL catalase, and 0.067 mg/mL glucose oxidase). The inset show the run length mean values normalized to control. While tau-mediated inhibition of kinesin-1 is reduced by phosphomimetic Y18E tau, the inhibition on kinesin-3 by tau increases with Y18E tau, as depicted on the schematic.

### Kinesin-3 enrichment in the periphery excludes tau

We next examined the localization of kinesin-3 (KIF1A) in COS-7 cells (Fig. 6). In cells expressing KIF1A alone (Fig. 6A) or with low tau expression, KIF1A localizes dynamically on peripheral microtubules (Fig. 6B). When tau is present, KIF1A enriches in stable protrusions. KIF1A also enriches on curved sections of microtubules in cells co-expressing tau (Fig. 6C, insets), reminiscent of tau enrichment on curved microtubules (Balabanian et al., 2017; Samsonov et al., 2004). Interestingly, we found that when KIF1A and tau (WT tau or tau Y18E) were co-transfected in cells, tau is excluded from binding KIF1A-enriched microtubules in the periphery (Fig. 6C), whereas tau binds all microtubules when expressed alone (Fig. S2A, B). These results support the idea that KIF1A and tau compete for access to the microtubule lattice (Lessard et al., 2019).

**Figure 6:**
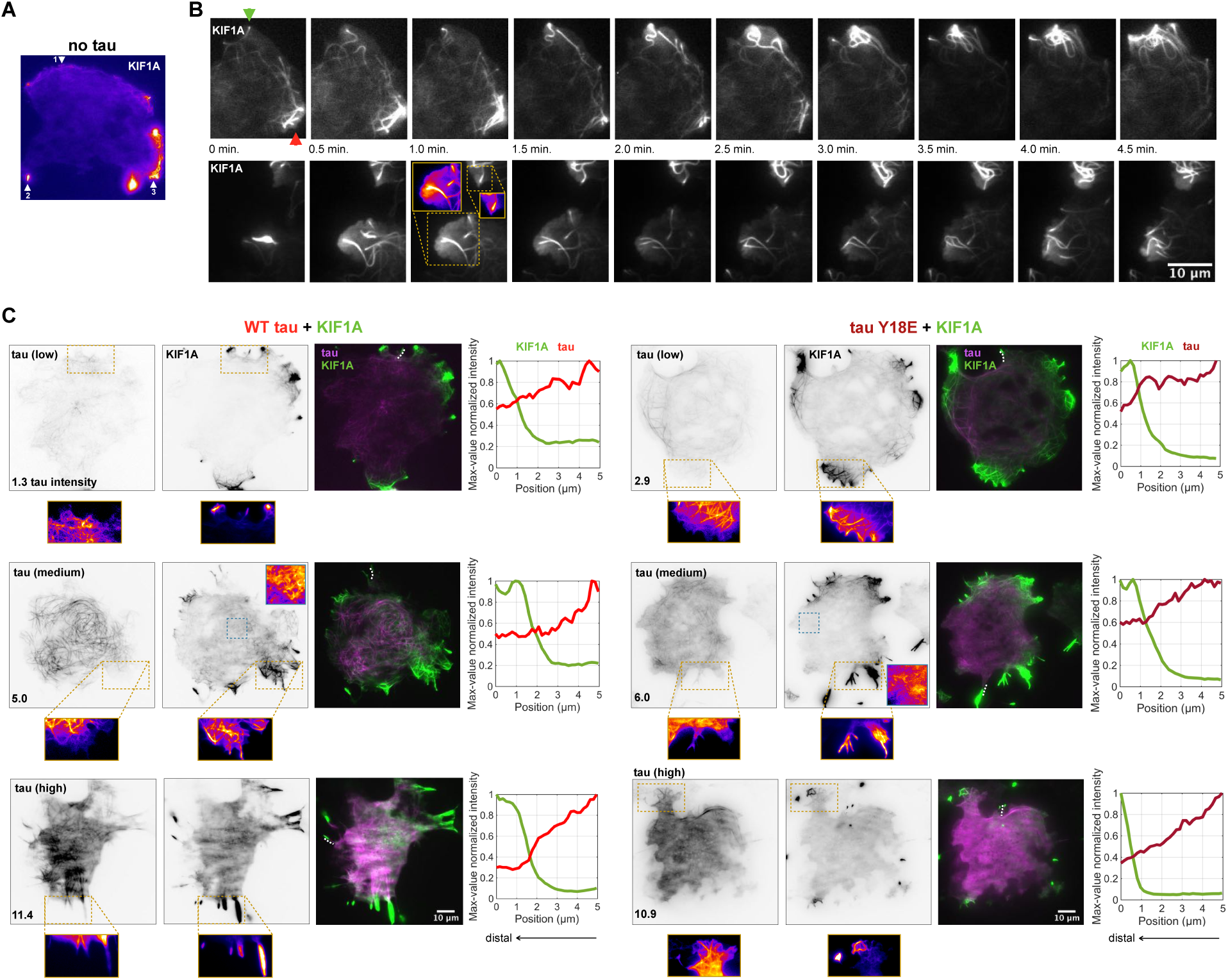
KIF1A localizes to peripheral microtubules and excludes tau binding. **A**. In cells transfected with KIF1A, KIF1A dynamically localizes at microtubule tips (arrow 1), accumulates on microtubules in protrusions (arrow 2), and binds along the length of peripheral dynamic microtubules (arrow 3) (Movie S4). **B**. The dynamic localization of KIF1A is also observed in cells expressing low levels of tau (normalized WT tau intensity here is 1.2 for top panels and normalized Y18E tau intensity is 1.7 for bottom panels), and microtubules exhibit polymerization dynamics. In the example here, KIF1A binds a growing microtubule tip, which probes close to the membrane (green arrow) and the KIF1A intensity increases at the microtubule tip. More KIF1A-bound microtubule ends grow and converge into that area with time. KIF1A intensity decreases gradually below (red arrow). KIF1A intensities also change in cytosolic areas around KIF1A-enriched dynamic microtubules (insets), while the membrane protrudes outward. **C**. KIF1A localization in the periphery excludes WT and phosphomimetic tau microtubule binding. The merged images show little overlap between tau and KIF1A. Intensity values of KIF1A and tau normalized to their respective maximum intensity value, over the regions of interest (white dotted line on merged images) show that tau intensity decreases as KIF1A signal rises. Additionally, we observe that KIF1A enriches on curved microtubules in cells co-expressing tau (insets). (KIF1A without tau: 6, WT tau + KIF1A: 40, Y18E tau + KIF1A: 45 observed cells.)

## Discussion

Intracellular cargoes navigate a complex cytoskeleton to their specific destinations in the cell. Tight control of vesicular trafficking is required to maintain signalling and degradative pathways. By organizing microtubules and altering the interaction between motor proteins and their microtubule tracks, microtubule-associated proteins are a key component of the regulatory network. Through high-resolution tracking and spatiotemporal motility analysis of vesicles, we show that tau differentially regulates early endosomes and lysosomes. Early endosome motility is inhibited by even low levels of tau, while lysosome motility is less sensitive. Early endosomes are inhibited by both unphosphorylated (WT) tau and phosphomimetic (Y18E) tau, while lysosome motility is not strongly affected by phosphomimetic tau. Taken together, our results indicate that tau has specific effects on the motility of different cargoes.

When cells over-expressed WT tau, the processivity of early endosomes and lysosomes is reduced, with fewer cargoes undergoing directed motion while lysosomes are less sensitive to phosphomimetic tau. In agreement, phosphomimetic mutations of the longest tau isoform in the central nervous system at 18 serine/threonine sites, many of which are normally phosphorylated by GSK-3 kinase, did not significantly affect the kinetics of moving tau particles driven by motor proteins, but rather increased the number of motile particles in axons, compared to WT tau (Rodriguez-Martin et al., 2013). Our results also indicate that unphosphorylated (WT) tau reduces the probability of cargoes to undertake directed motion, whereas phosphorylation of tau relieves this inhibition for specific cargoes. Phosphorylation is thought to reduce tau inhibition on cargo transport by introducing negative charges on tau, which might attract the positively charged motor domains of kinesin (Tarhan et al., 2013). Additional negative charges on tau might also reduce the affinity of tau to the microtubule surface, increasing the diffusivity of tau and in turn, decreasing the probability of hindering kinesin’s microtubule attachment (Rodriguez-Martin et al., 2013; Stern et al., 2017). The negative charge at Y18 also possibly alters the conformation of tau to favour the dynamic state, reducing the likelihood of tau acting as an obstacle to kinesin and allowing kinesin to efficiently navigate the microtubule surface (Stern et al., 2017).

The variation in responses of different cargoes to tau is likely determined by the types of kinesin motors driving their movement. While characterization of the set of motor proteins and adaptors that transport different cargoes is ongoing, current evidence suggests that kinesins 1, 2, and 3 contribute to early endosome and lysosome motility, but that the relative stoichiometry and activity of these kinesins varies among different cargoes. Kinesin-2 (KIF3A/B) is the primary anterograde motor for lysosomes (Brown et al., 2005). The more persistent motility of lysosomes on tau-decorated microtubules (Fig. 7), relative to early endosome motility, might be explained by the typically greater processivity of lysosomes in normal conditions and the presence of kinesin-2, which is known to be less sensitive to tau compared to kinesin-1 and kinesin-3. Kinesin-2 has a longer and more flexible neck-linker which is thought to help it circumvent tau molecules on the microtubule surface (Chaudhary et al., 2018; Dixit et al., 2008; Hoeprich et al., 2014; Monroy et al., 2020; Vershinin et al., 2007). Dynein motility is largely insensitive to tau (Chaudhary et al., 2018; Dixit et al., 2008; Vershinin et al., 2008), and we also found that retrograde motility of lysosomes is not inhibited by tau (Fig. 3D). For bidirectional cargoes driven by both kinesins and dyneins, tau reduces the number of engaged kinesins, biasing movement to the microtubule minus end (Chaudhary et al., 2018). In agreement, tau increases the fraction of inward-directed motility of lysosomes (Fig. 1D, E), resulting in clustering of the stationary lysosomes near the cell center (Fig. 1A, Fig. S4D).

**Figure 7:**
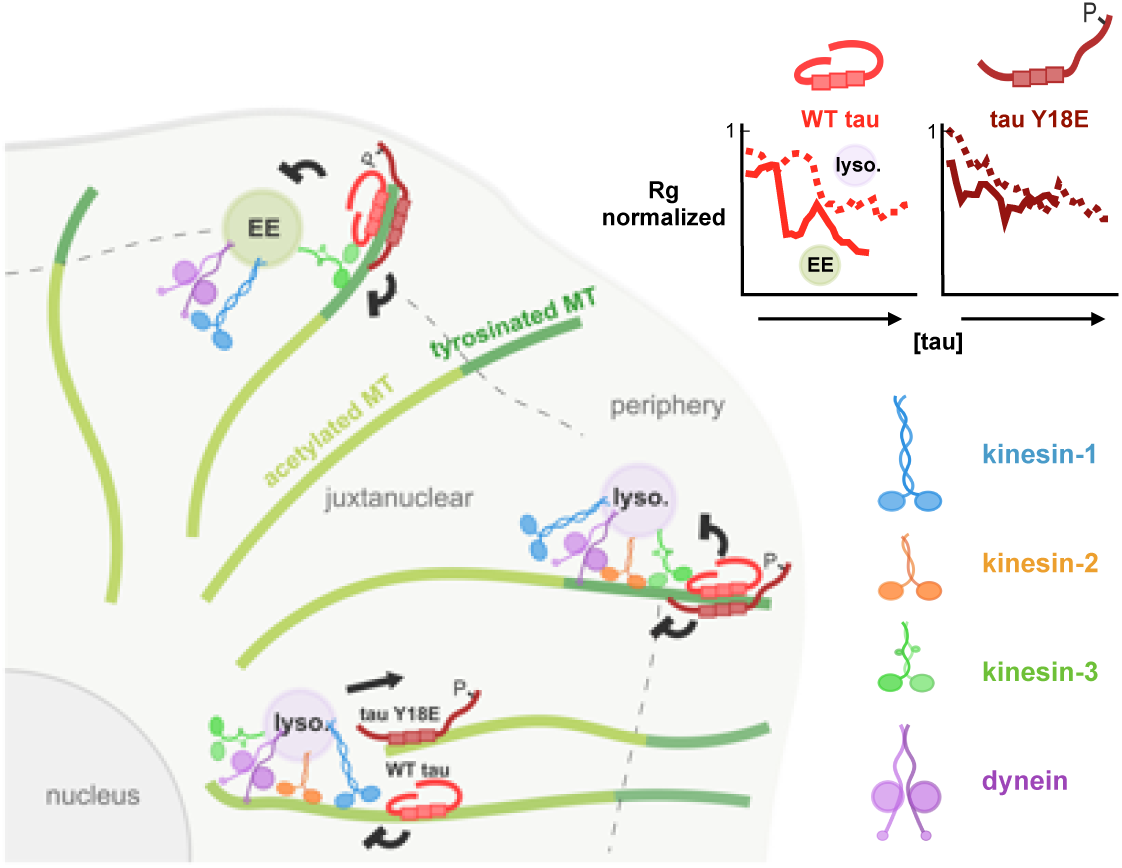
Model for the regulation of vesicle transport by tau Y18 phosphorylation. Kinesin-1 driven lysosome (lyso.) transport is inhibited by WT tau but is less sensitive to phosphomimetic tau. Kinesin-3 driven transport of early endosomes (EE) and peripheral lysosomes is inhibited by both WT and phosphomimetic tau. The Radius of gyration (Rg) (normalized to control) of early endosomes shows a steeper decrease with increasing tau levels in cells indicating that early endosome motility is more sensitive to tau inhibition compared to lysosomes.

When phosphomimetic mutations are made to tau (Y18E), inhibition of kinesin-1 is relieved *in vitro* (Stern et al., 2017), while kinesin-3 is more strongly inhibited (Fig. 5). In support of the motors specific to each cargo mediating differential regulation by tau, peripheral lysosomes and early endosomes are driven by kinesin-3 (Bentley et al., 2015; Guardia et al., 2016; Lenz et al., 2006; Matsushita et al., 2004). We observe that these cargoes are strongly inhibited by WT tau and more so, by phosphomimetic tau (Fig. 3, 4). In contrast, lysosomes closer to the nucleus are expected to be localized on acetylated microtubules and driven by kinesin-1 (Cai et al., 2009; Guardia et al., 2016; Katrukha et al., 2017; Norris et al., 2014; Tas et al., 2017). These juxtanuclear lysosomes maintained levels of motility similar to control in the presence of phosphomimetic tau (Fig. 3). Our live cell and *in vitro* observations show that modifying tau by phosphorylation modulates its effect on cargo transport, and that the effect of tau phosphorylation is likely dependent on the engaged motor proteins specific to different endocytic cargoes (Fig. 7).

Kinesin-3 (KIF1A) interacts with the tubulin C-terminal tails, which prolongs its contact with the microtubule and causes temporary pausing of the motor protein. KIF1A can then undertake more consecutive runs following pauses (Lessard et al., 2019). As tau also interacts with the C-terminal tails of tubulin, important for its diffusive behaviour (Hinrichs et al., 2012), it is thought that kinesin-3 and tau compete for the same binding sites on the microtubule (Lessard et al., 2019)(Fig. 6C). Our results indicate that phosphomimetic tau at Y18 is more inhibitory to KIF1A *in vitro* than WT tau, possibly because the higher diffusivity of phosphomimetic tau (Stern et al., 2017) increases its potential occupancy of more tubulin C-terminal tails over a wider microtubule surface. In turn, this decreases the pausing behaviour of kinesin-3 due to limited access to tubulin C-terminal tails and thus its overall run lengths are reduced over long distances.

The cell directs cargo transport by dynamically modulating interactions between microtubule-associated proteins (MAPs) and the microtubule surface by post-translational modifications of microtubules and MAPs, based on cellular functions and development. For example, during neuronal differentiation, proximo-distal gradient of tau phosphorylation states are regulated dynamically, with differing levels of tau phosphorylation in the soma and proximal axon compared to the axonal growth cone (Mandell and Banker, 1996). We observed that controlling tau phosphorylation differentially mediates the motion of cargoes varying in function and positioning. The implication is that aberrant tau phosphorylation causes dysregulation of the trafficking and targeting of organelles to specific sites in the cell.

## Materials and Methods

### DNA constructs

Tau-mApple, consisting of the human 3RS-tau sequence, the shortest isoform expressed in the central nervous system (confirmed by BLAST alignment), fused at the C-terminus to mApple, was a gift from Michael Davidson (mApple-MAPTau-N-10, Addgene plasmid # 54925). Using the QuikChange II XL Site-Directed Mutagenesis kit (Agilent, Wilmington, DE), we generated the phosphomimetic Y18E of this tau construct, by changing the plasmid’s nucleotide sequence coding the amino acid at position 18 from TAC to GAA, from tyrosine (Y) to glutamic acid (E). We confirmed the nucleotide changes with Sanger Sequencing using the standard CMV-F primer, at the McGill University and Genome Quebec Innovation Centre (Montreal, QC). The primers (Invitrogen) designed and used for the site-directed mutagenesis are the following: 5’-C T G T C C C C C A A C C C T T C C G T C C C A G C G T G - 3 ‘ a n d 5 ‘ - CACGCTGGGACGGAAGGGTTGGGGGACAG-3’. pRK5 c-Fyn, consisting of the human brain isoform B of fyn (confirmed by BLAST alignment) was a gift from Filippo Giancotti (Addgene plasmid # 16032)(Mariotti et al., 2001). DNA plasmids were purified using anion-exchange gravity flow chromatography (Qiagen, Hilden, Germany).

### Cell culture and transfection

COS-7 simian kidney fibroblast cells (American Type Culture Collection, Manassas, VA) were passaged with 0.25% trypsin-EDTA (Gibco, Thermo Fisher Scientific, Waltham, MA) and cultured in MatTek glass-bottom dishes (No. 1.0 Coverslip)(MatTek Corporation, Ashland, MA) in DMEM (Gibco) supplemented with 10% (v/v) FBS (Gibco) and 1% (v/v) Glutamax (Gibco). Cells were incubated at 37°C with 5% CO_2_for 24 h prior transfection. Cells were then transfected with 400 ng of DNA plasmid prepared in OPTI-MEM Reduced Serum (Gibco), using Lipofectamine LTX with Plus-Reagent, according to the manufacturer’s instructions (Invitrogen, Thermo Fisher Scientific). The transfection media containing DNA-lipid complexes was removed from the dishes after 4 hours and the cells were incubated at 37°C with fresh complete DMEM media overnight (∼12 h), before imaging.

### Live cell lysosome imaging

Cells were incubated with 50 nM Lysotracker Deep Red (Invitrogen) for 10 minutes in complete DMEM media. Cells were then washed and imaged in Leibovitz’s L-15 Medium (no phenol red) (Gibco) with 10% (v/v) FBS, at 37°C, using a custom Total Internal Reflection Fluorescence (TIRF) set-up built on an Eclipse Ti-E inverted microscope (Nikon, Melville, NY) attached to an EMCCD camera (iXon U897, Andor Technology, South Windsor, CT). Time-lapse movies were taken at 300 ms per frame for live cell imaging, with a 1.49 numerical aperture oil-immersion 100x objective (Nikon, Melville, NY) using argon ion Spectra Physics lasers (1 mW)(MKS Instruments, Andover, MA). The pixel size was 0.160 μm. Cells were imaged within 1 hour following Lysotracker staining.

### Endocytosis of Qdots and imaging

To image early endosomes, 0.02 μM streptavidin-conjugated Qdot 525 (Invitrogen) were coated with 2.4 μg/mL biotin-conjugated Epidermal Growth Factor (EGF)(Invitrogen) in Qdot Incubation Buffer (50 mM borate buffer, pH 8.3, 2% BSA)(Invitrogen) by incubating for 30 minutes on ice (Zajac et al., 2013). Cells were incubated on an orbital shaker with a solution of 2.2 μL Qdot preparation in a total of 200 μL complete DMEM media for 10 minutes. Cells were then washed and incubated with 1x CellMask (Invitrogen) complete DMEM solution for 5 min at 37°C. Finally, cells were washed and imaged in Leibovitz’s L-15 (supplemented with 10% FBS) before 1 hour post-internalization (Loubery et al., 2008; Scott et al., 2004) of Qdots (time_0_= start of incubation of cells with Qdots). Qdots were imaged at 37°C with the same instrumentation and parameters as lysosomes.

### Quantification of tau intensity

The expression of tau-mApple following transfection varied considerably from cell to cell. To reduce variability for data analysis and to investigate the roles of tau in a dose-dependent manner, expression levels of tau were quantified based on the intensity of tau’s mApple fluorescence signal in ImageJ (National Institutes of Health, Bethesda, MD). The intensities were normalized by the background intensity (area away from the cell) in the following manner: normalized tau intensity = cell-wide tau mean intensity/ background mean intensity. Experimental and imaging conditions were consistent within and across all experiments. Cells were categorized in 3 levels of tau expression: low, medium and high. In all cases, tau bound the microtubule filaments. Cells with low levels of tau expression (normalized tau intensity < 5) showed tau localizing nearly exclusively on microtubules with negligible levels of unbound cytosolic tau. At medium levels of tau (5 ≤ normalized tau intensity < 10), cells exhibited moderate amounts of cytosolic tau signal, mostly excluded from the peripheral region. Finally, cells with high levels of tau (normalized tau intensity ≥ 10) showed a high presence of tau in the entire cytosol.

### Retrospective staining of fyn kinase and microtubules

Following live cell imaging of lysosomes, cells were fixed with 4% (w/v) paraformaldehyde (Sigma-Aldrich, St. Louis, MO) for 10 min. Cells were then washed with Blocking Buffer: 2% (w/v) BSA (BioShop Canada Inc., Burlington, ON), 0.2% (w/v) saponin (Sigma-Aldrich) in PBS 1x (HyClone, Cytiva, Marlborough, MA). Fyn (15) primary antibody (Santa Cruz Biotechnology, Dallas, TX) was used to retrospectively stain for fyn kinase, α-tubulin primary antibody (Sigma-Aldrich) was used for total tubulin staining, and acetyl-α-tubulin primary antibody (clone 6-11B-1, Invitrogen) for acetylated tubulin. Following 1 h staining with primary antibody, cells were incubated with Alexa Fluor labelled secondary antibodies (Invitrogen) for 30 min., and imaged in PBS 1x with TIRF microscopy. Control cells show punctate signal with fyn antibody, while cells over-expressing fyn show cell-wide high signal (Fig. S1A).

### High resolution tracking of cargoes

Time-lapse movies for lysosomes and early endosomes were corrected for photobleaching using the histogram matching algorithm in ImageJ (Miura et al., 2014). Organelles were tracked with TrackMate (Tinevez et al., 2016), using the Laplacian of Gaussian filter to detect cargoes with sub-pixel localization and the simple Linear Assignment Problem (LAP) tracker for generating trajectories. The tracking parameters assume a maximal velocity of cargoes of 5 μm/s. Trajectories were imported into MATLAB (MathWorks, Natick, MA) to filter them and determine their directionality and localization (generated from polar plots), as well as their radius of gyration (Rg) and MSD. Only organelles detected in the first frame of movies and present for at least 5 frames were included in the analysis.

### Tracking-based analysis of lysosome directionality

Lysosome trajectories (from 90 seconds movies) were projected on polar plots with the MTOC designated as the cell center (0, 0 x, y coordinates). Moving lysosomes (Rg ≥ 0.5 μm) were parsed as having a net outward or inward directionality, based on the rho value of the first and last points in their trajectories. The percentages of stationary, outward or inward moving lysosomes are the mean of cell means with SEM.

### STICS analysis of lysosome directionality

Spatio-temporal image correlation spectroscopy (STICS) correlates fluorescence intensity fluctuations of image series in space and time to measure velocities and angles of mobile molecules and complexes. STICS combines the spatial information measured by 2D spatial correlations, with the time-dependent transport measured by temporal correlations. This allows us to define a spatio-temporal correlation function (CF), which is a function of the spatial lags (ξ,η) and the time lag (*Δt*)(Hebert et al., 2005):

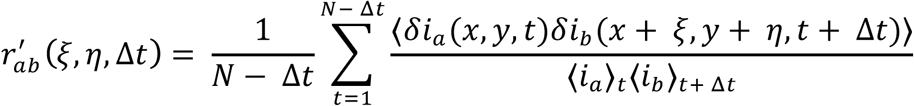

N is the total number of images in the time-series, r_ab_’ represents the average cross-correlation function for channels *a* and *b*, for all pairs of images separated by a time lag. We applied STICS on lysosome movies to determine directionality of motion. The temporal region of interest (TOI) size was 5 frames (1.5 s), corresponding to the mean directional switching time for lysosomes. The first 30 frames were analyzed for each movie. The ROI size was set to 16 × 16 pixels. We manually established the coordinates of the cell center and calculated the angle of movement with respect to the cell center (see Fig. 1E for details) to determine the direction of lysosomes. Vector angles of 0-60 degrees represent inward motion towards cell center, 60-120 degrees as perpendicular motion and 120-180 degrees as outward motion.

### Analysis of lysosomal localization

The rho value of a trajectory on polar plots is the mean distance (or mean rho) of the trajectory’s points (from each time frame) from the center. The rho value was normalized to the distance from cell center to cell edge, which was estimated by measuring the cell area using the formula: radius = √(cell area/π). Lysosomes were considered to be localized peripherally when normalized rho of trajectory ≥ 0.85, in the juxtanuclear region when 0.5 ≤ rho < 0.85 and in the perinuclear region when < 0.5 (method used in Fig. 3B-E, Fig. S4D-F). The percentages of lysosomes categorized based on directionality represent the mean of cell means with SEM.

Alternatively, the number of peripheral lysosomes was counted using the Laplacian of Gaussian filter detection algorithm of TrackMate (Tinevez et al., 2016), and divided by the total number of detected lysosomes in the cell. The periphery was defined as ∼6 μm from the cell edge. Following the tracing of the cell contour, the inner peripheral line was drawn using the eroding feature in ImageJ. Cells with a total area > 2000 μm^2^were selected for this analysis, as the perinuclear cloud of smaller cells is typically very close to the periphery (method used for Fig. S4A, C).

### Motion analysis of cargoes

The radius of gyration (Rg) of lysosome or early endosome trajectories (from 90 seconds movies) was calculated with the following equation:

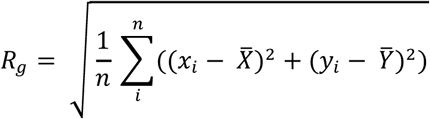

Each trajectory consists of a number *n* of detected spots at consecutive time frames. Here, the Rg of a trajectory represents the deviation in 2D space of the position *x*_*i*_ and *y*_*i*_ of a time point *i* within the trajectory from the mean position of all spots, 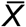 and 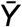. Rg is the average deviation from the center of mass and thus can be simply thought as the radius of a circle that contains half of the spots of the trajectory. In other words, the Rg is a measure of the travelled distances of lysosomes or early endosomes from the center of their trajectory. Minimal thresholds for Rg were not applied when studying the motion of all lysosomes or early endosomes (Fig. 2, Fig. 4). A minimal Rg threshold of 0.5 μm was used to differentiate between stationary and moving lysosomes (Fig. 1, Fig. 3).

The mean squared displacement (MSD) of particle trajectories in × and y (2 dimensions) was analyzed using a package (MSDanalyzer) developed for MATLAB (Tarantino et al., 2014). The first 10% of log-log transformed MSD curve over 90 seconds movies (for lysosomes) or 30 seconds movies (early endosomes) were fitted to a linear function to determine the exponent α value (Tarantino et al., 2014). Fits of low quality (R^2^coefficient < 0.8) were discarded from the distribution of α values. The standard error of the mean was calculated as SEM ≈ weighted standard deviation /√number of degrees of freedom (Tarantino et al., 2014). Overall, the trends from the Rg and MSD measurements are in agreement.

### Microtubule density analysis

Microtubule organization was studied by qualitatively studying the network configuration. Microtubules are typically organized radially from the MTOC, with few microtubules bending or bundling in the periphery; this type of organization was considered normal (Fig. S2E top schematic). With higher levels of tau expression, many microtubule filaments were seen circling the periphery (Fig. S2E left schematic) or forming dense bundles in a ring-like configuration (Fig. S2E right schematic). In addition, the peripheral density of the microtubule network was analyzed by calculating the mean intensity of tubulin staining in the peripheral region (∼6 μm from the cell edge) normalized to the mean background intensity.

### *in vitro* KIF1A motility assays

KIF1A(1-393)-LZ-3xmCitrine motors (a gift from Dr. Kristen Verhey, University of Michigan, Ann Arbor, MI) were expressed in COS-7 monkey kidney fibroblasts (American Type Culture Collection, Mansses, VA) as previously described (Lessard et al., 2019). 3RS-tau expression, mutagenesis, purification, and labelling were conducted as previously described (Stern et al., 2017). Bovine tubulin isolation and paclitaxel-stabilized, fluorescently labelled microtubule polymerization were conducted as previous described (Lessard et al., 2019). In experiments containing tau, microtubules were polymerized in the absence of labelled tubulin. Microtubules were then incubated with 200 nM of Alexa 647 labelled 3RS-tau for 20 minutes at 37ºC. After incubation, microtubules were centrifuged at room temperature for 30 minutes at 15,000 rpm and resuspended in motility buffer. KIF1A *in vitro* motility assays were conducted as previously described (Lessard et al., 2019). Of note, *in vitro* motility assays were conducted in a motility buffer of 12 mM PIPES, 1 mM MgCl_2_, 1 mM EGTA, supplemented with 20 μM paclitaxel, 10 mM DTT, 1 mM MgCl_2_, 10 mg/ml BSA, 2 mM ATP and an oxygen scavenger system [5.8 mg/ml glucose, 0.045 mg/ml catalase, and 0.067 mg/ml glucose oxidase]). Motility events were analyzed as previously reported (Hoeprich et al., 2017; Hoeprich et al., 2014). In brief, overall run length and speed data was measured using the ImageJ (v. 2.0.0, National Institute of Health, Bethesda, MD) MTrackJ plug-in, for a frame-by-frame quantification of KIF1A motility. Run length and standard deviation were calculated and adjusted for microtubule track length as previously reported (Thompson et al., 2013).

### Tau immunostaining of iPSC-derived neurons and COS-7

Human induced pluripotent stem cells (iPSC) from the line AIW-002-02 (Montreal Neurological Institute and Hospital, Montreal, QC) were cultured in mTESR plus medium (STEMCELL Technologies Inc., Vancouver, BC), on Matrigel (Corning, New York, NY) coated dishes at 37°C with 5% CO_2_. Briefly, monolayer iPSC cultures were differentiated into neural progenitor cells (NPCs) with STEMdiff Neural Induction Medium, and NPCs were differentiated into neurons using BrainPhys hPSC Neuron Kit, following the manufacturer’s instructions (STEMCELL Technologies Inc.). After 10 days, neurons were washed with PBS 1x pre-warmed at 37°C and fixed with 4% paraformaldehyde for 15 minutes. Coverslips were rinsed 3x with PBS 1x, permeabilized with 0.3% Triton X-100 (Sigma-Aldrich) for 5 min. and then incubated with Blocking Buffer (2% BSA, 0.1% Triton X-100 in PBS 1x) for 1 hour at room temperature. Neurons were incubated with tau-5 (clone 5) primary antibody (Sigma-Aldrich) for 2 hours, and with Alexa Fluor 647 labelled secondary antibody (Invitrogen) for 45 min., following washes with Blocking Buffer. Coverslips were mounted on slides with ProLong Diamond Antifade Mountant (Invitrogen) and imaged with TIRF microscopy at 300 ms per frame exposure, on a 100x objective (Nikon).

COS-7 cells transfected with WT tau alone or with fyn kinase were similarly stained with tau-5 (Sigma-Aldrich) or with tau-9g3 primary antibody (GeneTex Inc., Irvine, CA) followed by Alexa Fluor 647 labelled secondary antibody (Invitrogen) at the same concentrations and incubation times as tau-5 staining of iPSC-derived neurons, and imaged with the same parameters.

### Statistical analysis

The sliding means of Rg and α values of lysosome or early endosome trajectories with respect to tau intensity were averaged with a bin size of 5 along the x-axis, using bootstrapping over 1000 iterations and with 95% confidence intervals to determine statistical significance (Chaubet et al., 2020). The sliding means of cell data points were similarly calculated, except that the fraction of the total data points included in individual bins and the number of bins averaged for the means were adapted based on sample size, but kept constant across conditions per analysis. The distribution of α values were analyzed using 1-way ANOVA and multiple comparison post-hoc Tukey’s test for comparing control, low levels of WT tau and Y18E tau. 1-way ANOVA and Tukey’s test were used to determine differences between the distributions of stationary or moving lysosome percentages in the peripheral, juxtanuclear and perinuclear regions of cells expressing low levels of WT tau and Y18E tau compared to control. Rg of moving lysosomes (in peripheral, juxtanuclear, perinuclear regions) were log-normalized and 1-way ANOVA with Tukey’s test were used to determine the p-values between control, low levels of WT tau and Y18E tau conditions. We also confirmed the results with the Kruskal-Wallis non-parametric test (without log-normalization). 2-way ANOVA (2×2 factorial design) was used to distinguish the effects of low levels of WT tau, fyn, and the interaction of WT tau with fyn on vesicle motility in cells co-expressing WT tau and fyn. For *in vitro* motility assays, effects on overall run lengths and overall speed between control, WT tau and Y18E tau were determined using 1-way ANOVA and post-hoc Tukey tests. All statistical tests were performed with MATLAB (MathWorks).

## Supplemental material

The supplemental material consists of 4 movies and 5 figures. Fig. S1 includes the lysosome directionality analysis for cells expressing low, medium or high tau levels, the analysis for cells co-expressing WT tau and fyn, and the results of the tau antibody stainings in COS-7 cells and iPSC-derived neurons. Fig. S2 presents the analysis of microtubule density. Fig. S3 supports the lysosome motility analysis and presents the pausing analysis. Fig. S4 presents additional methodology used to measure the percentages of peripheral lysosomes, and presents additional data to distinguish the different responses of lysosome distribution, motility and directionality in subcellular localizations. Fig. S5 supports the motility analysis of early endosomes. Movie S1 and Movie S2 show the motility of lysosomes and Qdot-containing early endosomes respectively in cells. Movie S3 shows the motility of purified KIF1A motor proteins on reconstituted microtubules. Movie S4 shows the dynamic localization of KIF1A in a COS-7 cell.

## Acknowledgements

The authors thank Loïc Chaubet for adapting the computation of bootstrap means to our data, Emily Prowse for preparing coverslips with differentiated iPSC-derived neurons and Abdullah R. Chaudhary for important feedback and discussions on the analysis. We thank the McGill University and Genome Quebec Innovation Centre for sequencing the phosphomimetic Y18E tau-mApple plasmid.

This work was supported by the National Institutes of Health (NIH) grant R01 GM132646-02 to C.L. Berger and A.G. Hendricks, the Canadian Institutes of Health Research (CIHR) grant PJT-159490 to A.G. Hendricks and the Natural Sciences and Engineering Research Council of Canada (NSERC) grant RGPIN-2017-05005 to P. W. Wiseman.

The authors declare no competing financial interests.

## Author contributions

L. Balabanian and A.G. Hendricks designed the cellular assays, and L. Balabanian performed the preparations, experiments and analysis. P. Yaninska, P.W. Stevens, and P.W. Wiseman performed the STICS analysis, D.V. Lessard and C.L. Berger designed the *in vitro* motility assay experiments and D.V. Lessard performed the experiments and analyzed the data. L. Balabanian prepared the plots and figures with feedback from A.G. Hendricks. L. Balabanian and A.G. Hendricks wrote the manuscript. D.V. Lessard and C.L. Berger wrote the Methods section on *in vitro* motility assays and P. Yaninska and P.W. Wiseman wrote the Methods section on STICS. All authors edited the manuscript.

## SUPPLEMENTAL MATERIAL

**Movie S1:** TIRF time-lapse movie of lysosomes stained with Lysotracker in a control COS-7 cell (Fig. S1A). The video was captured at 300 ms per frame, corrected for photobleaching (see Methods). 9x real-time.

**Movie S2:** TIRF time-lapse movie of internalized streptavidin-conjugated Qdot 525 coated with biotin-EGF in a control COS-7 cell (Fig. S5A). The video was captured at 300 ms per frame, corrected for photobleaching (see Methods). The background was subtracted only for the purpose of presentation (rolling ball radius of 15 pixels in ImageJ). 9x real-time.

**Movie S3:** TIRF single-molecule motility of KIF1A(1-393)-LZ-3xmCitrine on reconstituted microtubules *in vitro* without tau (control)(Fig. 5A). The video was captured at 200 ms per frame. 6x real-time.

**Movie S4:** TIRF time-lapse movie of KIF1A in COS-7 cell transfected with KIF1A only (Fig. 6A). The video was captured at 300 ms per frame. 27x real-time.

**Figure S1:**
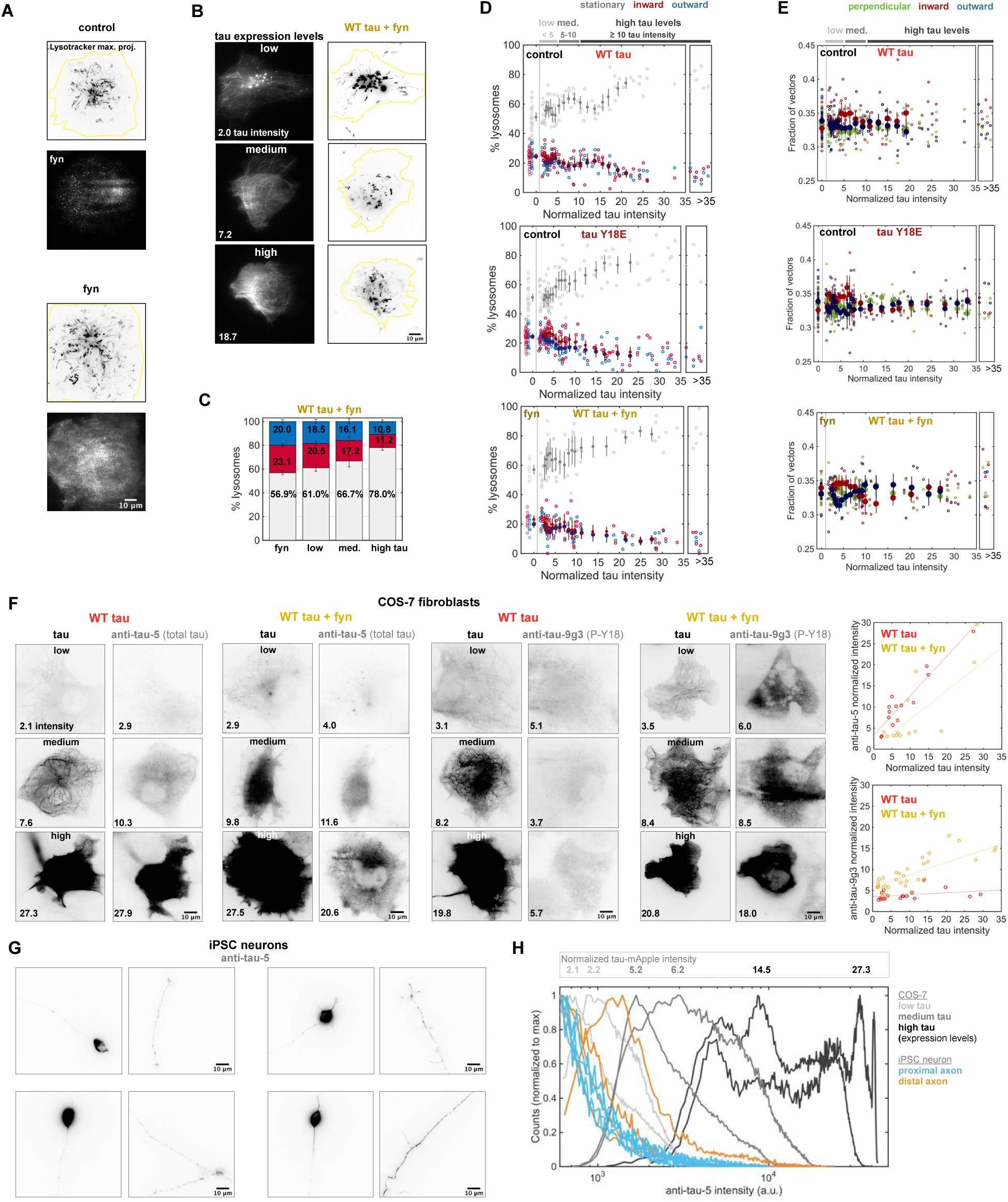
High tau levels strongly inhibit lysosome motility. Endogenous tau levels in iPSC neurons is consistent with low expression in COS-7 cells. **A**. Maximum projection of a control cell (Movie S1) shows robust lysosome motility. Fyn kinase overexpression resulted in strong cell-wide signal from immunostaining, as opposed to the sparse puncta observed in control. **B**. Cells were co-transfected with WT tau and fyn kinase, which phosphorylates tau at Y18. Tau levels were quantified as low, medium or high. **C**. The percentage of stationary lysosomes increases in fyn over-expressing cells, compared to control (see Fig. 1C), and with tau co-expression. **D**. The percentages of inward and outward moving lysosomes of cells as a function of tau expression (low, medium and high) quantified using single-particle tracking, and **E**. spatiotemporal image correlation spectroscopy (STICS) demonstrate a bias of inward motion of lysosomes with tau (mean ± 95% confidence intervals, using bootstrapping). The trend is more apparent in cells expressing low levels of tau, as at high tau expression levels, fewer lysosomes are moving and microtubule organization is altered (Fig. S2). (Control: n = 35 cells in D and 33 in E; fyn: 11; WT tau: 71; Y18E tau: 83; WT tau + fyn: 50 cells.) **F**. WT tau was transfected in cells alone or with fyn kinase, and the intensity tau-mApple was measured to determine tau expression levels (x-axis in the plot on the righthand side). Cells were then fixed and stained with primary tau-5 antibody (with Alexa 647 secondary antibody) to measure the total tau levels (y-axis in the plot), or with tau-9g3 antibody to measure the levels of tau phosphorylated at Y18. Tau-5 antibody staining intensity increases with tau levels in cells expressing WT tau alone or co-expressed with fyn kinase. Tau-9g3 staining intensity increases with tau levels in cells co-expressing WT tau and fyn kinase, indicating that fyn kinase phosphorylates tau at Y18. Tau-9g3 staining intensity does not increase with tau levels in cells expressing WT tau alone, without fyn kinase overexpression. (WT tau, anti-tau-5: n = 14 cells; WT tau + fyn, anti-tau-5: 13; WT tau, anti-tau-9g3: 18; WT tau + fyn, anti-tau-9g3: 31.) **G**. Tau is enriched in the distal axon of fixed iPSC-derived neurons, at day 10 post terminal differentiation (BrainPhys Neuronal Medium (STEMCELL Technologies Inc.)). **H**. We used identical immunofluorescence protocols to stain for total tau (tau-5 antibody) and quantify fluorescence intensity in iPSC neurons and COS-7 cells. Intensities in axons are similar to low tau expression levels in COS-7 cells. Tau levels are higher in the distal axon than in the proximal axon.

**Figure S2:**
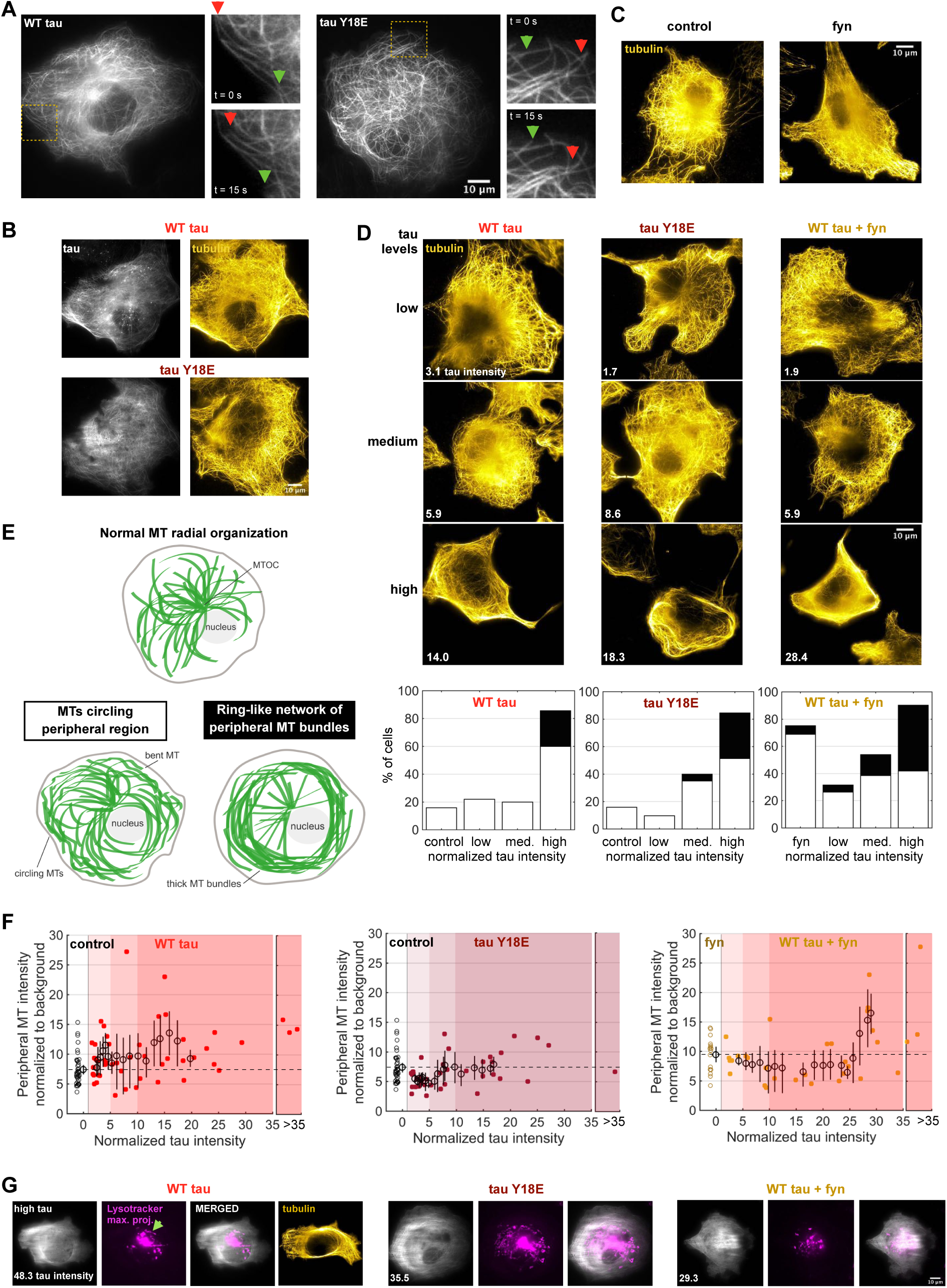
Tau increases microtubule density and bundling in the cell periphery. **A**. mApple-tau decorates microtubules. Microtubule dynamics persist in cells expressing low levels of WT tau or Y18E tau, where microtubules assemble (green arrows) and depolymerize (red arrows). **B**. When expressed in fibroblasts, tau binds along the length of microtubules. Slight differences in the tau and tubulin images stem from the time between imaging tau in live cells, and then fixing and staining for microtubules. **C, D**. Following live cell imaging, cells were fixed and stained for microtubules. High levels of tau increase peripheral microtubule density and rearrange the microtubule network into thick ring-shaped bundles (lower panels). **E**. Microtubules are normally arranged radially from the MTOC (microtubule organizing center) with few microtubules circling in the peripheral region (top schematic). In some cells expressing low levels of tau, bent peripheral microtubules were seen circling near the plasma membrane (left schematic). This was relatively more common for WT tau cells than tau Y18E, in agreement with the quantification of microtubule density in F. With high levels of tau expression, more cells show high microtubule density and substantial circling in the periphery (left schematic, white bars), or a ring-shaped bundled microtubule network (right schematic, black bars). The majority of control cells and cells with low levels of WT tau or Y18E tau maintain a normal radial microtubule organization. (Control: n = 38 cells; WT tau: low 32, medium 10, high 35; Y18E tau: low 31, medium 20, high 39; fyn: 16; WT tau + fyn: low 19, medium 13, high 31 cells.) **F**. Peripheral microtubule density (∼6 μm from cell edge) was quantified from the tubulin immunostaining intensity of cells and normalized to background away from the cell. The sliding means of the peripheral microtubule intensity of cells and 95% confidence intervals (open circles with bars) were calculated using bootstrapping. The dashed lines represent the control or fyn means. (Control: n = 39 cells; WT tau: 48; Y18E tau: 38; fyn: 16; WT tau + fyn: 32 cells.) **G**. Lysosome localization is affected by the microtubule network’s rearrangement at very high levels of tau expression where lysosomes are trapped within the microtubule bundled arrays (3rd schematic in E), and some are seen circling the interior edges (green arrow).

**Figure S3:**
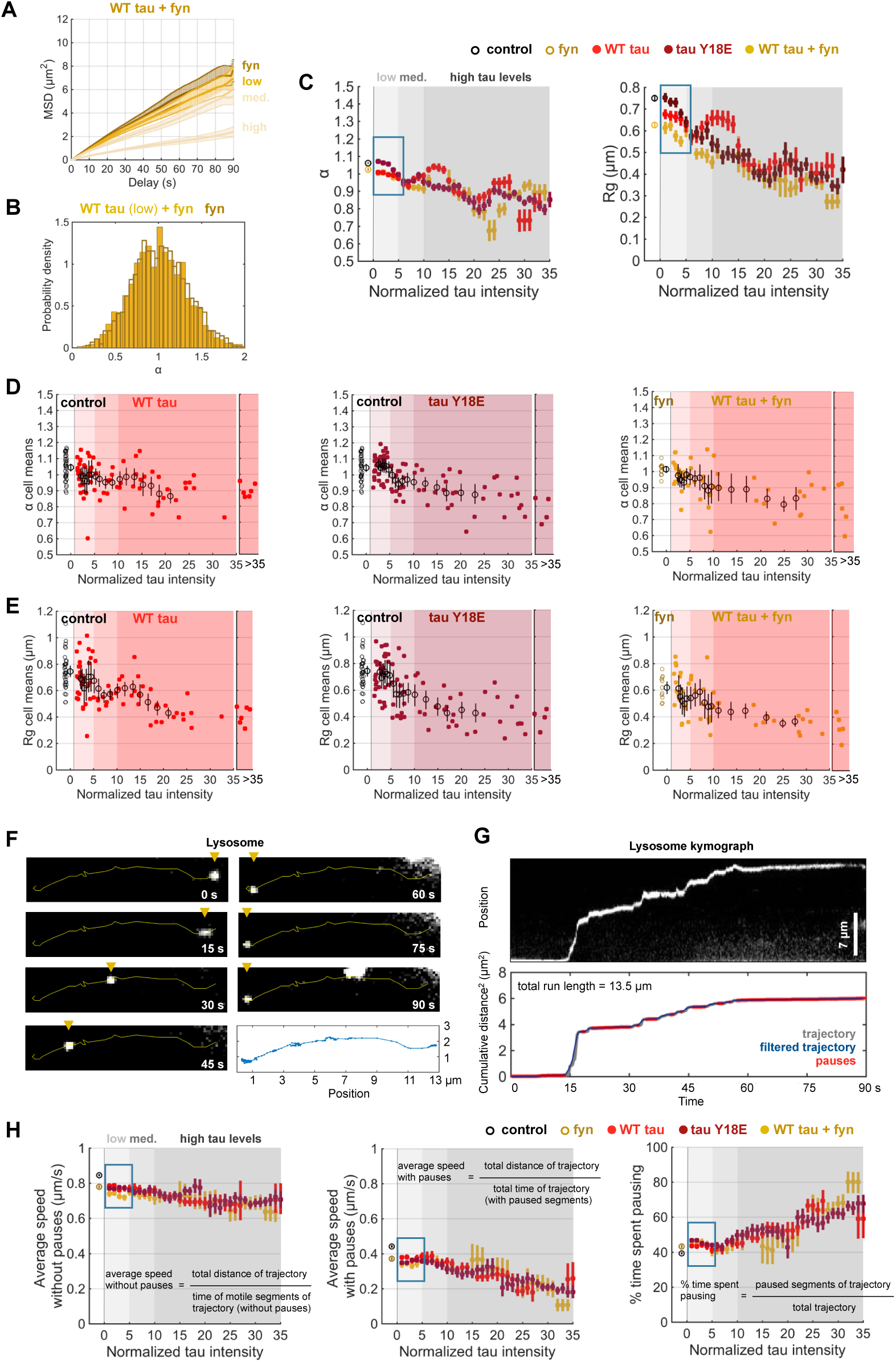
Tau inhibits lysosome motility and increases pausing. **A**. MSD and **B**. the distribution of α values of lysosome trajectories shows that lysosome motility in cells co-expressing low levels of WT tau and fyn kinase is similar to that of cells over-expressing fyn only (2-way ANOVA, tau effect: p < 0.0001, fyn effect: p = 0.0008, interaction tau and fyn: p = 0.013). **C**. The α and Rg means of lysosome trajectories decrease as a function of tau expression levels. Low levels of tau (blue rectangles) co-expression with fyn do not further inhibit lysosome motility (95% confidence intervals), compared to fyn expression alone. The plots showing the means for cells expressing WT tau and tau Y18E, normalized to control are found in Fig. 2D, E. **D**. The mean of MSD α values and **E**. Rg (radius of gyration) for each cell analyzed as a function of tau expression levels (determined by tau intensity normalized to the background) show similar trends compared to plots pooling all trajectories across cells within conditions as seen in C. (Control: n = 35 cells; WT tau: 72; Y18E tau: 83; fyn: 11; WT tau + fyn: 50 cells.) **F**. Time-lapse of moving lysosome is compared to the trajectory rendered by tracking with Trackmate (last panel). **G**. The kymograph from the movie of the same lysosome is compared to the kymograph generated with tracking (below). Paused segments in the trajectory (in red) are defined by consecutive frame distances of less than 0.100 μm and correspond to pauses observed on the kymograph above. **H**. The sliding means of the average speed of moving lysosome trajectories (Rg ≥ 0.5 μm) not including the detected paused segments decreases slightly with increasing tau levels. The average speeds of trajectories including the paused segments show a stronger reduction with tau levels, explained by the increased pausing times with tau (mean ± 95% confidence intervals). (Control: n = 5408 trajectories, from 35 cells, over 20 experiments; fyn: 2421 traj., 11 cells, over 7 experiments; WT tau: 9024 traj., 73 cells, over 14 experiments; Y18E tau: 11105 traj., 83 cells, over 5 experiments; WT tau + fyn: 5379 traj., 50 cells, over 8 experiments.)

**Figure S4:**
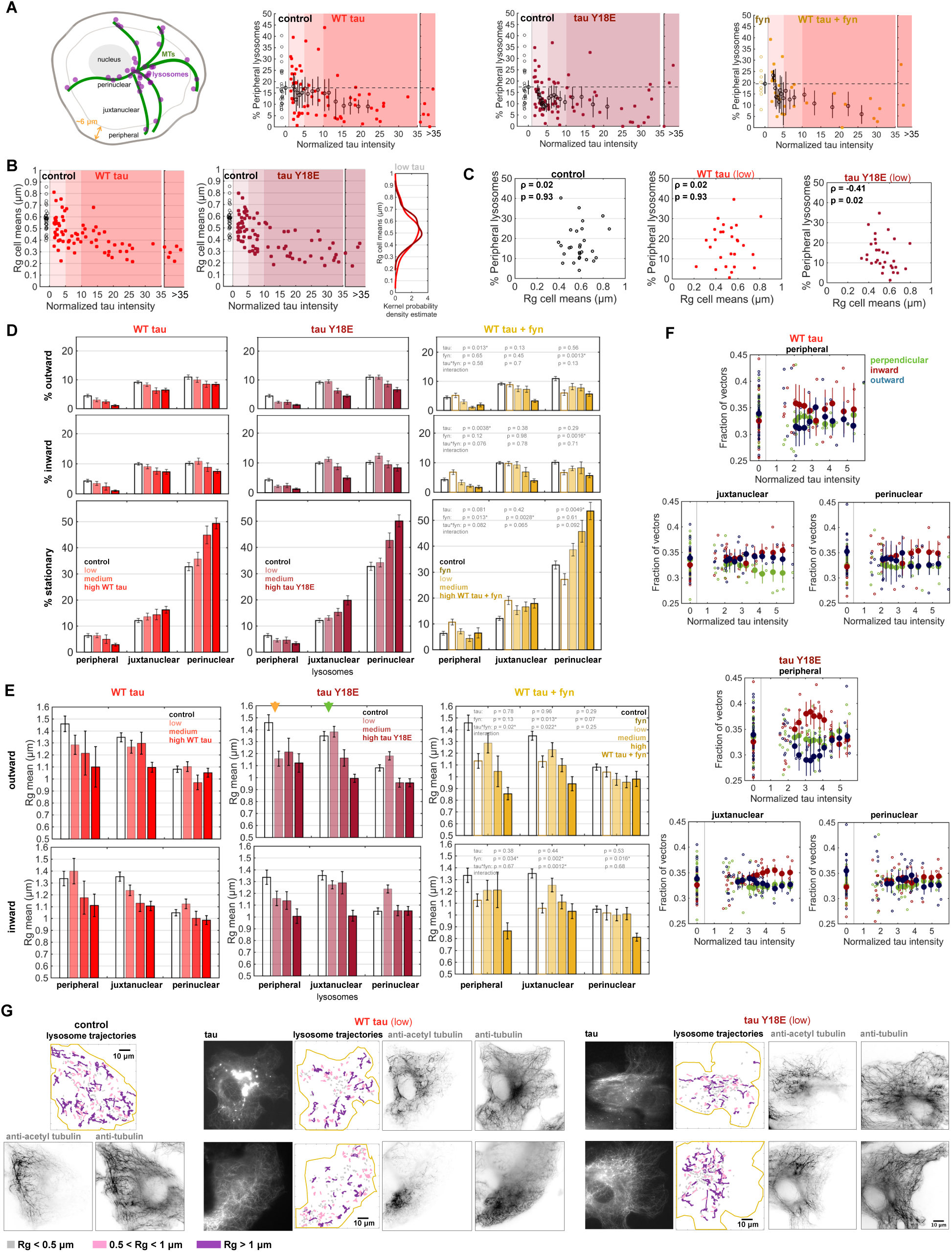
Peripheral lysosomes are strongly inhibited by tau Y18E. **A**. We define the peripheral region of the cell as ∼6 μm from the cell edge. The percentage of peripheral lysosomes is defined as the number of lysosomes in the peripheral region over all detected lysosomes with Trackmate. The percentage of peripheral lysosomes decreases with increasing tau levels (small symbols indicate individual cells, large symbols indicate mean ± 95% confidence intervals calculated using bootstrapping). Cells expressing low levels of Y18E tau have a lower percentage of peripheral lysosomes compared to control and cells expressing low levels of WT tau. Cells expressing fyn show a higher percentage of lysosomes in the periphery compared to control, in agreement with D. Small cells (cell area < 2000 μm^2^) were excluded from the analysis. (Control: n = 28 cells; fyn: 8; WT tau: 68; Y18E tau: 75; WT tau + fyn: 29 cells.) **B**. The mean Rg for the lysosome trajectories in each cell analyzed (over 30 sec. movies) decreases monotonically as a function of WT tau levels. The Rg of Y18E tau expressing cells suggest two subpopulations, one with average Rg ∼ 0.6 μm and another with average Rg ∼ 0.5 μm. Probability density estimate based on normal kernel function for the distribution of Rg cell means for cells expressing low levels of WT tau shows one peak, whereas Y18E tau shows two peaks, suggesting bimodality of lysosome movement (right plot). A low Rg cell mean was considered as Rg < 0.5 μm (over 30 sec. movie) and high as > 0.6 μm (examples shown in Fig. 3A). (Control: 35 cells; WT tau: 72; Y18E tau: 83 cells.) **C**. The percentage of peripheral lysosomes were defined here with the method used in A. Control cells and WT tau expressing cells show no correlation between the percentage of peripheral lysosomes and Rg mean of lysosome trajectories (over 30 sec. movies). Y18E tau expressing cells with lower percentage of peripheral lysosomes have higher Rg cell means (significant moderate inverse correlation, Spearman’s correlation coefficient ρ = -0.41, p-value = 0.02). If the mean of the Rg of lysosome trajectories in a Y18E tau expressing cell was low (Rg < 0.5 μm, in A), constrained lysosomes were found in the peripheral region (Fig. 3A top panels). If the Rg mean of a cell was high (Rg > 0.6 μm, in A), lysosomes were largely absent from the peripheral region (Fig. 3A bottom panels). In both cases, most long runs were positioned centrally, in the juxtanuclear region. Together, these observations suggest that peripheral and juxtanuclear lysosomes are differentially affected by phosphomimetic Y18E tau. (Control: 28 cells; WT tau (low): 27; Y18E tau (low): 31 cells.) **D**. The plots here are extensions of those shown in Fig. 3C, to include the data from cells not only expressing low levels of tau, but also medium and high, as well as from cells co-expressing WT tau and fyn. Fyn reduces the proportion of moving lysosomes in the perinuclear region, while increasing the presence of stationary lysosomes in the periphery (2-way ANOVA for cells co-expressing WT tau (low) and fyn, p-values on plots for effects of tau, fyn, and interaction of tau and fyn). More stationary lysosomes are constrained in the perinuclear region with increasing tau levels. **E**. Similarly, the plots here are extensions of those shown in Fig. 3D. Fyn inhibits lysosome motility in both directions (2-way ANOVA for cells co-expressing WT tau (low) and fyn, p-values on plots). **F**. STICS analysis indicates that WT and more significantly Y18E tau preferentially alter lysosome directionality in the cell periphery, where tau expression increases the fraction of inward motility (mean ± 95% confidence intervals). **G**. Following lysosome imaging, cells were fixed and stained for acetylated microtubules and all microtubules. Lysosome trajectories are colour-coded based on Rg. Lysosomes with longer runs are localized in peripheral and central regions, in regions with acetylated and non-acetylated microtubules, of control cells and cells expressing WT tau. However, long trajectories are limited to the central regions, which coincide with acetylated microtubules, in cells expressing low levels of Y18E tau. Lysosomes are largely absent in the peripheral regions of Y18E tau expressing cells.

**Figure S5:**
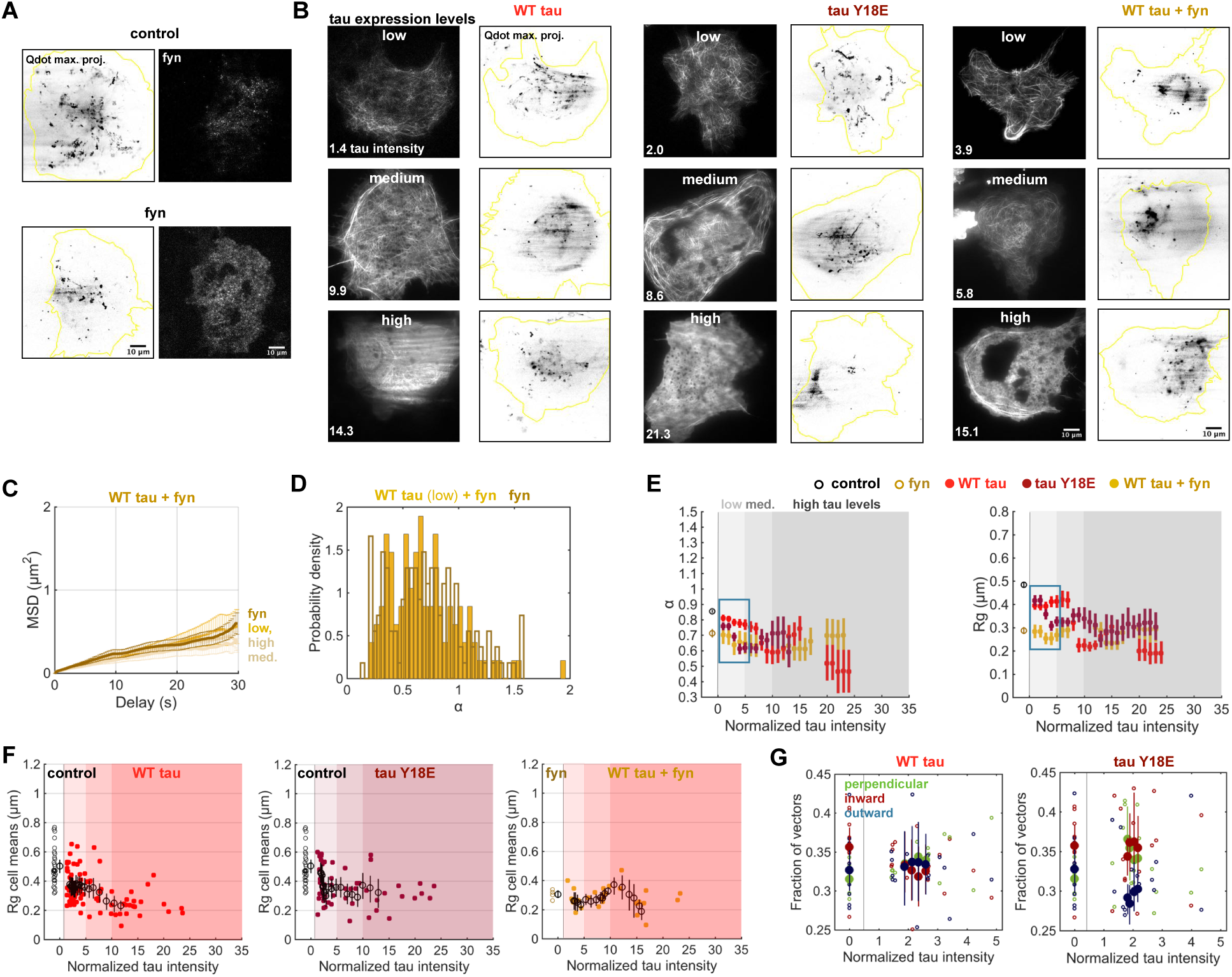
WT tau and phosphomimetic tau inhibit early endosome motility. **A**. Maximum projection of early endosome motility in a control cell (Movie S2) indicates that early endosomes move relatively smaller distances than lysosomes. **B**. The maximum projections show that early endosome motility is strongly inhibited by tau, particularly at medium and high levels of tau. **C**. MSD analysis and **D**. α values of early endosome trajectories show that fyn expression strongly reduces early endosome processivity, such that further inhibition by low levels of tau is not discernible with fyn (2-way ANOVA, tau effect: p = 0.24, fyn effect: p < 0.0001, interaction tau and fyn: p = 0.51). **E**. The cell means of α and Rg (radius of gyration) of early endosome trajectories decrease with increased levels of tau expression (mean ± 95% confidence intervals, see Fig. 4D, E). **F**. The displacement (Rg) of early endosomes is inhibited strongly by tau, even at low levels of tau expression (small symbols indicate mean values for individual cells, large open symbols indicate sliding mean and 95% confidence intervals). The trends here are similar compared to plots pooling all trajectories across cells within conditions (like in E). **G**. STICS analysis demonstrates that Y18E tau perturbs outward movement (mean ± 95% confidence intervals). (Control: n = 1946 trajectories, from 28 cells, over 19 experiments; fyn: 268 traj., 4 cells, over 3 experiments; WT tau: 2783 traj., 84 cells, over 15 experiments; Y18E tau: 2193 traj., 53 cells, over 6 experiments; WT tau + fyn: 687 trajectories, 26 cells,over 6 experiments.)

## Notes

### Competing Interest Statement

The authors have declared no competing interest.

